# Real-time imaging of rotation during synthesis by the replisome

**DOI:** 10.1101/2025.04.01.646591

**Authors:** Thomas M Retzer, Lional T Rajappa, Masateru Takahashi, Samir M Hamdan, Karl E Duderstadt

## Abstract

During chromosome replication, unwinding by the helicase and synthesis by the polymerases can lead to overwinding and supercoiling of DNA. The mechanical consequences of these events and resulting local dynamics at the replication fork are not well understood. To address these issues, we developed a transverse DNA flow-stretching approach to spatially resolve the parental, leading and lagging strands in real-time. Using bacteriophage T7 as a model system, this approach revealed bursts of high-speed replisome rotation that support continuous DNA synthesis. Surprisingly, excessive rotation does not reduce replisome speed, but increases pausing, reduces processivity, and increases the number of polymerases. Taken together, our observations reveal intrinsic pathways to overcome challenges posed by unfavorable DNA topologies during DNA replication.

## Introduction

Throughout the domains of life, DNA is copied using a similar mechanism by replisomes that share a conserved core architecture (Yao and O’Donnell, 2016). Unwinding of parental DNA by a helicase is coupled to the synthesis of two daughter strands by DNA polymerases. Due to the antiparallel arrangement of DNA, the leading strand is synthesized continuously whereas the lagging strand is synthesized discontinuously as Okazaki fragments supported by repeated primer synthesis by primases. Coordination of these and many other moving components involved in chromosome replication is frequently challenged by obstacles such as DNA damage (Sparks et al., 2019), chromosome organizational elements (Gruszka et al., 2020) and other structurally diverse barriers (Zeman and Cimprich, 2014). Among these are chromosomal regions with unexpected DNA topologies (Postow et al., 2001; Guo et al., 2021) that challenge unwinding by the helicase. The mechanical consequences and resulting dynamics of the replisome during collisions with topological barriers have been experimentally challenging to address. As a consequence, it has remained unknown whether replisomes possess intrinsic pathways to cope with topological challenges to avoid genome instability. The double-helical structure of DNA is advantageous for the storage and maintenance of genetic information, but it poses major challenges when the information-rich DNA bases must be accessed during genome duplication. Unwinding by the helicase leads to overwinding in the parental DNA (Fig. 1A). Likewise, superhelical tension forms on the daughter strands as a consequence of DNA synthesis by the polymerases (Ullsperger et al., 1995) (Fig. 1B). To avoid chromosome damage and genome instability (Branzei and Foiani, 2010; Pommier et al., 2016; Adolph and Cortez, 2024), these unfavorable DNA topologies must be resolved by topoisomerases, which are a class of enzymes that help ensure chromosomes are maintained during genome compaction, chromosome segregation, DNA replication and transcription (Branzei and Foiani, 2010; Schoeffler and Berger, 2008; Koster et al., 2010; Pommier et al., 2022). Nevertheless, there are many regions on chromosomes where topoisomerases are unable to prevent the accumulation of overwinding. In particular, topoisomerases struggle to keep pace in highly transcribed regions, at topological boundaries, near chromosome ends and with rapidly moving replication forks (Stracy et al., 2019; Postow et al., 1999; Keszthelyi et al., 2016; Bermudez et al., 2010; Naughton et al., 2013; Achar et al., 2020; Lang and Merrikh, 2021). Replisome rotation has been proposed as an alternative mechanism to cope with overwinding resulting from helicase activity (Champoux and Been, 1980). The observation of interwound daughter strands and the formation of pre-catenines provides support for this proposal (Peter et al., 1998; Schalbetter et al., 2015; Snapka et al., 1988; Sundin and Varshavsky, 1981; Cebrian et al., 2015), but rotational motion of the replisome during synthesis has not been directly observed.

**Figure 1.**
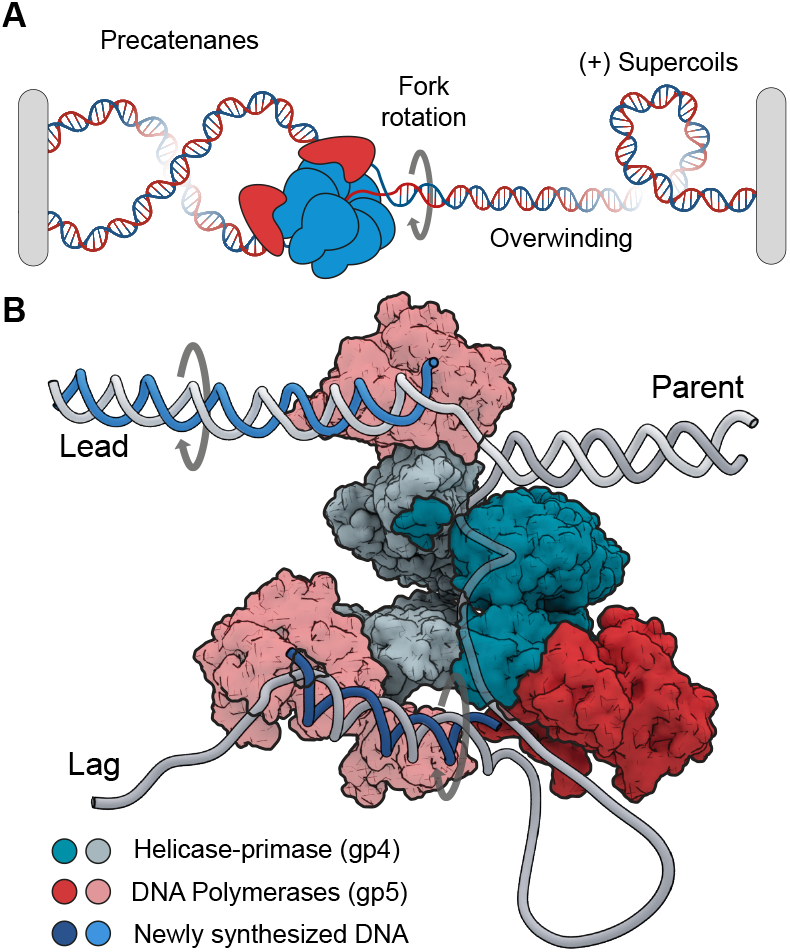
Topological challenges during DNA replication. **(A)** DNA replication introduces overwinding and positive supercoils. This can lead to fork rotation and the formation of precatenanes on the daughter strands. **(B)** Structural model of the T7 replisome (Gao et al., 2019). Parent DNA is unwound by the helicase generating templates (Lead and Lag) used for daughterstrand synthesis. The DNA polymerases travel on a helical path leading to rotation of the daughter strands or the formation of superhelical tension.

To address these issues, we used single-molecule fluorescence imaging to directly visualize replisome dynamics during topological challenges. We reconstituted the replisome from bacteriophage T7, which performs DNA replication with a minimal set of components (Hamdan et al., 2009; Gao et al., 2019). We quantified replication fork progression on DNA molecules with topological barriers introduced in the parental strand. Surprisingly, we find that the presence of barriers resulted in only a very modest reduction in replisome speed, however, this reduction was accompanied by an increase in pausing frequency and reduced processivity. To characterize the fundamental mechanics of these encounters, we developed a transverse DNA flow-stretching approach that allows for real-time spatial resolution of the parental, leading and lagging strands. Strikingly, this approach revealed that high speed replisome rotation supports bursts of continuous DNA synthesis during encounters with topological barriers. While allowing the replisome to overcome topological barriers, rotation does result in more polymerases residing at the replication fork, consistent with frequent disruptions to helicase-polymerase coordination. Taken together, our observations reveal replisome rotation is an intrinsic pathway that supports continued DNA synthesis during topological challenges, but at the cost of detrimental changes in replisome coordination.

## Results

### Visualization of DNA replication during encounters with topological barriers

We developed a total internal reflection fluorescence (TIRF)-based single-molecule approach to visualize the dynamics of DNA replication during encounters with topological barriers. DNA replication was reconstituted using purified components in a stepwise manner. First, 27-kb-long DNA molecules were surface-immobilized on functionalized coverslips via biotin-streptavidin-biotin interactions (fig. S1A). A replication fork was introduced at one end of the molecules with a single biotin on the leading-strand for surface attachment and one or multiple biotins for surface attachment of the opposite end. We created two types of molecules: unconstrained (using a single biotin and allowing the parental strand to rotate freely) and constrained (using multiple biotins and creating a topological barrier that prevented free rotation of the parental strand) (Fig. 2A). In the case of a single biotin, the parental strand was free to rotate (unconstrained), whereas, in the other case, multiple biotins created a topological barrier in the parental strand that prevented free rotation (constrained) (Fig. 2A). The purified T7 replication proteins gp5-trx (DNA polymerase), gp4 (helicase-primase), and gp2.5 (ss-DNA binding) were introduced to start the reaction (fig. S2A). Finally, flow through the flowcell was stopped and replication fork progression was imaged by fluorescently staining the DNA and monitoring accumulation of the lagging strand product, visible as a blob moving unidirectionally along the immobilized DNA molecules (Fig. 2B, fig. S2B).

**Figure 2.**
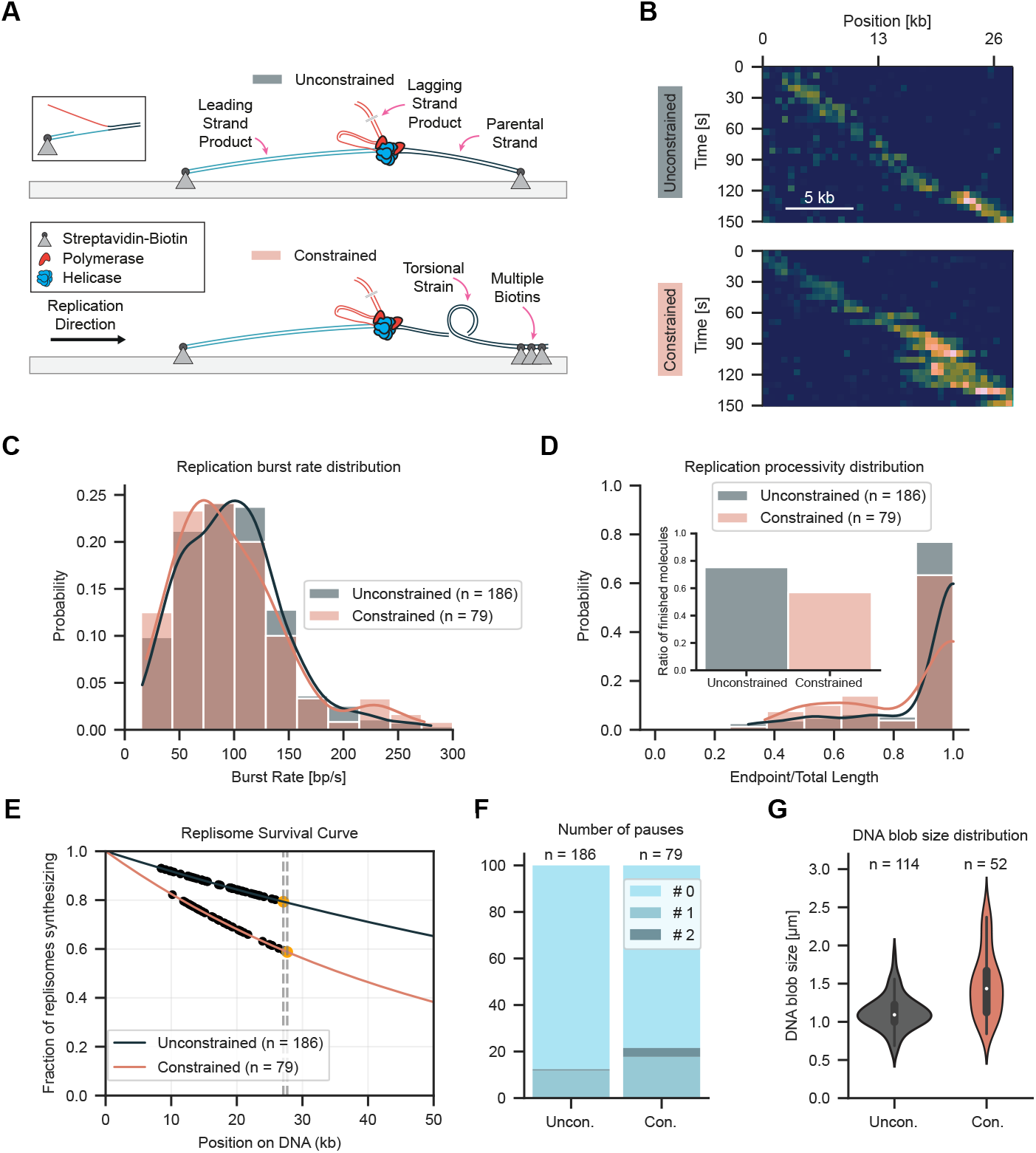
Visualizing replisome progression under topological strain. (**A**) Schematic of replication assay with unconstrained and constrained molecules. A preformed fork is used to initiate DNA replication using T7 replisome components. Leading and parental strands are tethered either by a single or multiple biotin interactions. Multiple biotin interactions ahead of the replisome create a topological challenge. (**B**) Representative kymograph showing DNA replication for unconstrained and constrained molecules. (**C**) Replication burst rate distribution for unconstrained and constrained molecules. (**D**) Replication processivity distribution for unconstrained and constrained molecules. Inset displays the fraction of molecules that replicated to the end. (**E**) Replication processivity estimation using a maximum likelihood estimation. (**F**) Pause probabilities for unconstrained and constrained molecules. (**G**) Replication blob size for unconstrained and constrained molecules.

The dynamics of replication on the unconstrained DNA were consistent with previous single-molecule observations. Replication kinetics were quantified by performing subpixel localization and tracking of the lagging-strand replication product (fig. S2C). This analysis resulted in a burst replication rate of 99 ± 3 bp/s (mean ± SEM, n

= 186 molecules) (Fig. 2C). A total of 75% of the unconstrained DNA molecules were completely copied resulting in 27 kb products (Fig. 2D). To accurately estimate the processivity, we used maximum likelihood estimation with a model that combined both replication events observed to stop before the end and those prematurely terminated when they reached the end. Using this approach, the estimated mean processivity was 117 kb (95% confidence interval from 102 to 137, n = 186 molecules) (Fig. 2E), higher than previous reports (Lee et al., 2006; Kulczyk et al., 2012; Duderstadt et al., 2016). Pausing was observed in 13 % of molecules with 12 % of replisomes pausing once, and 1 % pausing twice (Fig. 2F, fig. S3) at our experimental time resolution.

Several measures were taken to avoid photo-damage of DNA that causes nicks, disrupts DNA replication, and prohibits observations of DNA topology dynamics. Optimal conditions were established by quantifying the frequency of nicking using supercoiled DNA molecules attached at each end to the cover slip surface with multiple biotins as previously described (fig. S4) (Ganji et al., 2016). This approach allowed for the establishment of a low light intensity imaging condition in which 95 % of the DNA molecules remained intact during the observation time.

### Replisomes tolerate topological strain

Having established robust imaging conditions, next we performed replication with the constrained DNA. The presence of multiple biotin attachment sites at the end of the parental strand prohibiting free rotation would be expected to present a major challenge to helicase unwinding and DNA synthesis, which involves rotation along the helical axis of DNA. However, tracking the motion of the lagging-strand replication product (Fig. 2B, fig. S2) unexpectedly revealed a remarkable tolerance by the replisome when faced with the topological constraint. Surprisingly, the mean burst rate of 96 ± 5 bp/s (mean ± SEM, n = 79 molecules) was within the margin of uncertainty when compared to the unconstrained DNA (Fig. 2C). Nevertheless, the number of molecules that were replicated to completion was reduced to 57 %, with a mean processivity of 52 kb (95% confidence interval from 43 to 67, n = 79 molecules) (Fig. 2D, E). The frequency of molecules with pausing events increased to 22 % and there was an increase in the fraction of molecules exhibiting a second pause to 4 % (Fig. 2F, fig. S3). The observation of only modest reductions in processivity suggested the replisome must be exploiting an alternative pathway to relieve torsional strain. To test whether processivity could be restored by adding topoisomerase, we added a low concentration of DNA gyrase (5 nM) to the reaction. This resulted in a replication burst rate of 103 ± 5 bp/s (mean ± SEM, n = 97 molecules), which was within the margin of uncertainty compared with unconstrained and constrained DNA without gyrase (fig. S5A). The mean processivity increased to 84 (95% confidence interval from 70 to 105, n = 97 molecules), indicating a positive effect on replication at modest concentrations of DNA gyrase (fig. S5B, C). The frequency of pausing decreased to 10 %, with 9 % showing a single pause and 1 % showing two pauses (fig. S5D).

Visual inspection of kymographs of individual constrained molecules revealed elongation and shape variability in the lagging-strand products (Fig. 2B, fig. S2B). These shape changes were not observed in the unconstrained DNA where the lagging-strand product appears as a single uniform bundle moving in a unidirectional fashion. To quantify the behavior, the kymographs were segmented to determine estimates for the size of the lagging-strand product. Elongation was more pronounced once the products had increased in length toward the end of replication where a mean size difference from 1.1 µm (unconstrained, n = 114 molecules) to 1.5 µm (constrained, n = 52 molecules) was observed. Furthermore, size fluctuations doubled from a standard deviation of 0.2 µm (unconstrained) to 0.4 µm (constrained) (Fig. 2G). This pronounced shape variability hinted at a change in the mechanics of the replisome at the replication fork that could not be fully resolved.

### Transverse flow platform to study DNA topology dynamics

To further resolve the spatial dynamics of the lagging-strand products during DNA synthesis, we devised an alternative flowcell geometry in which replication could be monitored in the presence of a transverse flow. This was accomplished using an x-shaped flowcell with two perpendicular flow lanes that crossed each other at the location of imaging (Fig. 3A). DNA replication was reconstituted in a stepwise manner but with different flow lanes used for each step. DNA molecules were surface-immobilized on the functionalized coverslips using one flow lane. Next, the other flow lane was used to introduce the replication components using a flow transverse to the DNA molecules. In contrast to the assay presented in Fig. 2, where flow was stopped during imaging, and the lagging-strand product appeared as a compact blob, here transverse flow was continuously applied during imaging to extend the lagging-strand product during synthesis. With the new geometry, the lagging strand product appeared as a straight line that grew in length over time from the location of the moving replisome at the replication fork (Fig. 3B, fig. S6, Movie S1). Based on the direction of movement, molecules were segmented and the length of each arm of the replication fork was precisely measured (Fig. 3C). This approach spatially resolves the parental, leading, and lagging strands in real-time during DNA replication.

**Figure 3.**
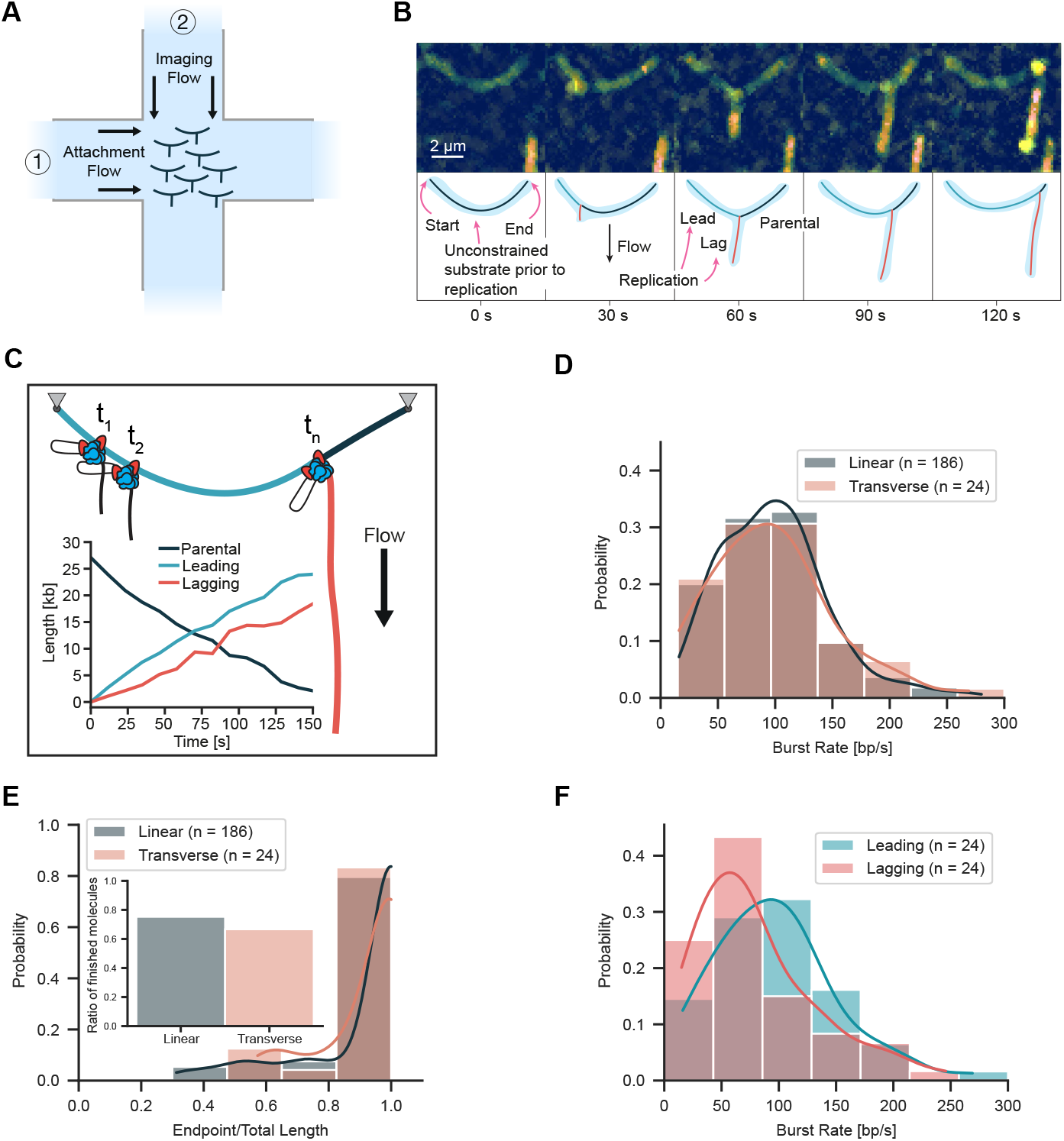
Visualization of DNA replication using transverse flow. (**A**) Schematic of transverse flow imaging of DNA replication. After surface immobilization of DNAs using the attachment flow, a perpendicular imaging flow is used to extend the lagging-strand products during DNA replication. (**B**) Representative kymograph of replication on an unconstrained DNA molecule that appears as an arch with the lagging-strand product extending from the arch over time. (**C**) Length of Parental, Leading, and Lagging strands as a function of time with a cartoon displaying strand organization. (**D**) Rate distribution for unconstrained molecules in the linear and transverse configurations. (**E**) Processivity distribution for unconstrained molecules in the linear and transverse configurations. Inset displays the fraction of molecules that replicated to the end. (**F**) Distribution of leading and lagging strand synthesis rates using based on the individual product lengths.

The application of transverse flow did not influence the replication kinetics of the unconstrained DNA. The mean burst rate was 97 ± 7 bp/s (mean ± SEM, n = 24 molecules) and the processivity was 84 kb (95% confidence interval from 60 to 140, n = 24 molecules) (Fig. 3D, E, fig. S7) with a confidence interval overlapping the processivity observed for the linear replication assay. The mean burst rate for the constrained DNA was 98 ± 9 bp/s (mean ± SEM, n = 28 molecules) (Fig. 4D), consistent with our findings in the absence of transverse flow. However, we observed a reduction in processivity for the constrained DNA to 29 kb (95% confidence interval from 21 to 46, n = 28 molecules) and the percentage of molecules fully replicated was reduced to 32 % (Fig. 4E).

**Figure 4.**
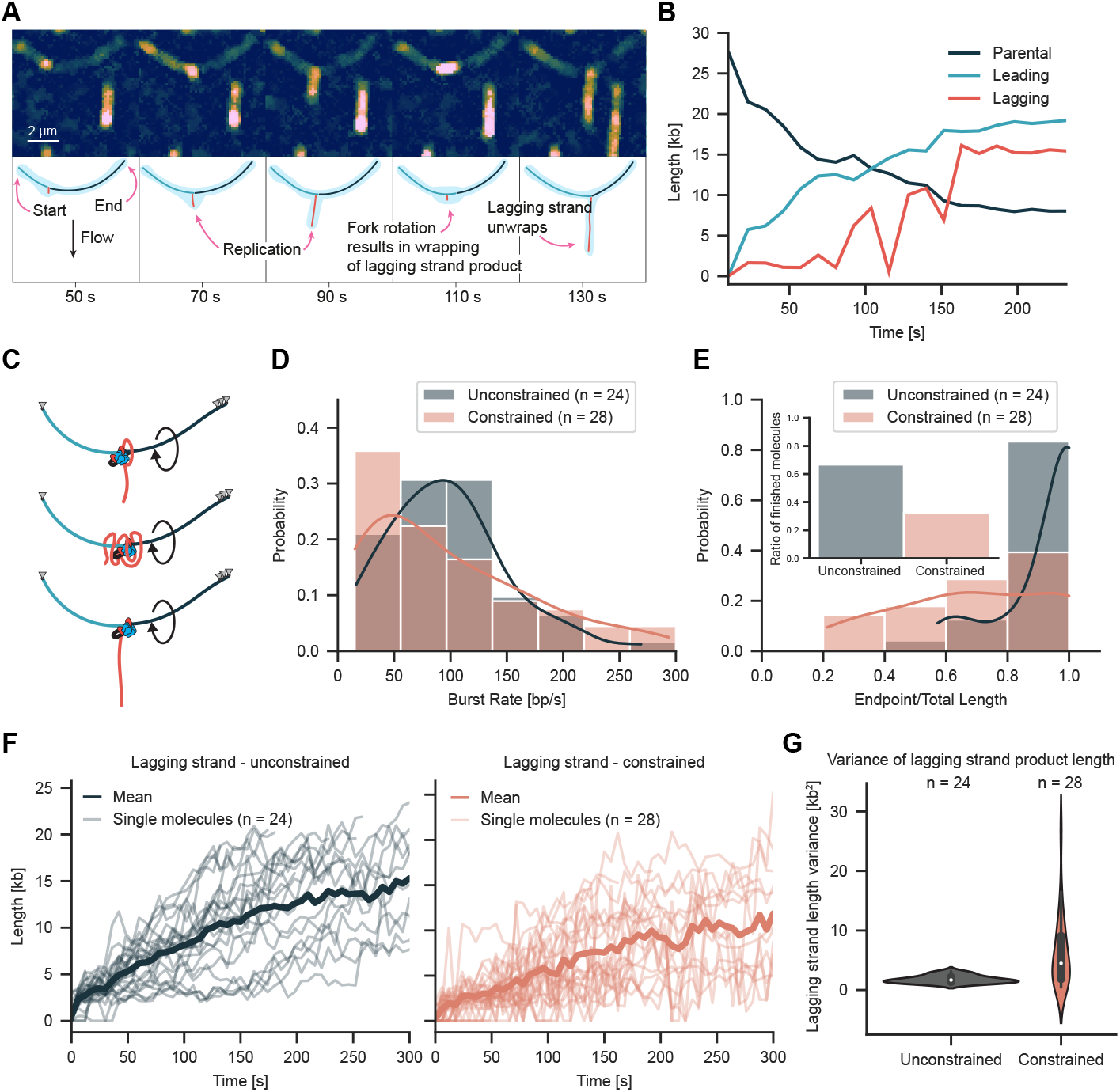
Transverse flow imaging reveals replication fork rotation. (**A**) Representative kymograph of replication on a constrained molecule that appears as an arch. The lagging-strand product appears as a blob wrapped along the arch. (**B**) Length of parental, leading, lagging strands over time for the molecule displayed in A. (**C**) Cartoon depicting mechanism of fork rotation and arch organization. (**D**) Rate distribution for unconstrained and constrained molecules using transverse flow imaging. (**E**) Processivity distribution for unconstrained and constrained molecules imaged using transverse flow. Insert bar graph shows fraction of completely replicated molecules. (**F**) Lagging strand length over time for unconstrained and constrained molecules. (**G**) Variance in lagging strand length for unconstrained and constrained molecules.

Transverse flow imaging spatially separates each of the strands emerging from the replication fork revealing the leading and lagging strand synthesis dynamics. Measurements of lagging-strand growth revealed a mean burst rate of around 80 ± 7 bp/s (mean ± SEM) which was lower than the leading-strand synthesis rate (Fig. 3F). We attribute this difference to the presence of compacted gp2.5-coated regions undergoing replication that are shorter in length than dsDNA at the applied force (Hamdan et al., 2009). Using length measurements of the individual strands of the replication fork, we estimated the applied forces with the worm-like chain model. The force on the leading and parental strands of the replication fork was 1.27 ± 0.59 pN (median ± MAD, n = 24 molecules) (fig. S8A). Force on the lagging strand is expected to increase over time as the lagging-strand product grows and the volume of DNA subjected to viscous drag from flow increases. We estimate a maximal force on the lagging strand of approximately 0.22 ± 0.05 pN (median ± MAD, n = 24 molecules) (fig. S8B) once a full product has been synthesized. Previous studies have shown that replication is not disrupted by forces in the low pN range used to make observations (Hamdan et al., 2009).

### Lagging strand dynamics reveal fork rotation

Transverse flow imaging of the constrained DNA revealed significant changes in replisome mechanics including bursts of replication fork rotation. For the unconstrained DNA, the lagging-strand product grew as a straight line over time. In contrast, significant length fluctuations were observed for the constrained DNA with the lagging-strand product frequently wrapping into a compact blob similar in structure to that observed in the linear replication experiments (Fig. 4A, fig. S9, Movie S2). Segmentation of the individual strands and tracking of the replisome position confirmed leading-strand synthesis continued during wrapping (Fig. 4B). These observations are consistent with rotation of the replication fork during synthesis and wrapping of the lagging-strand around the leading-strand (Fig. 4C). In cases where synthesis pauses and then resumes, the extended lagging-strand product adopts a blob consistent with rewrapping as fork rotation continues during synthesis (fig. S10). The burst replication rate of 98 ± 9 bp/s (mean ± SEM, n = 28 molecules) for constrained molecules was nearly identical to 97 ± 7 bp/s (mean ± SEM, n = 24 molecules) for unconstrained molecules (Fig. 4D). In contrast, we observed a 52% reduction in the number of molecules that were completely copied (Fig. 4E).

To quantify variation in the extension of the lagging strand product over time, length measurements were transformed into a stationary function by taking the difference of consecutive values (Fig. 4F, fig. S11). The mean difference between consecutive length measurements was more than three times greater for the constrained DNA, increasing from 1.8 ± 0.6 kb^2^ (mean ± SD, n = 24 molecules) for the unconstrained DNA to 6.1 ± 5.9 kb^2^ for the constrained DNA (mean ± SD, n = 26 molecules) (Fig. 4G). The larger standard deviation for constrained DNA is consistent with the significant length changes that are visible in the kymographs from individual molecules.

Topological barriers encountered during replication can lead to overwinding of the parental strand to accommodate the helical path of the replisome. At the low force applied by transverse flow, overwinding of the parental strand would be expected to lead to formation of positive supercoils (van Loenhout et al., 2012). However, we did not observe significant length variation in the parental strand or compaction of the arch structures formed by transverse flow. This indicated no positive supercoils formed on the parental strand during replication and instead only small transient changes in twist may have occurred. Taken together, these observations suggest fork rotation is the dominant pathway used by the replisome to overcome acute topological barriers on the parental strand under our experimental conditions.

The transverse flow applied to visualize lagging-strand synthesis and rotational dynamics favors a specific replication fork geometry. Under these conditions, we observe a reduction in processivity to 29 kb (95% confidence interval from 21 to 46, n = 28 molecules) for constrained molecules in comparison to 84 kb (95% confidence interval from 60 to 140, n = 24 molecules) for unconstrained molecules (fig. S7, Table S1). This represents a 65% reduction in processivity, which is greater than the 55% reduction observed for constrained vs unconstrained in the linear replication experiments. This difference may result from the specific replication fork geometry during transverse flow requiring additional torque to wrap the lagging-strand. Whether linear or transverse flow are more reflective of the replication fork geometry adopted in cells remains unclear. However, we imagine the replisome encounters significant forces when traveling through topologically challenging regions in the chromosome that could be similar if not more significant than the application of transverse flow. Notably, the motion of the lagging-strand is not restricted in our experiments, whereas in cells the presence of the daughter chromosome would be expected to constrain the motion.

### Fork rotation alters polymerase exchange dynamics

To determine if fork rotation leads to changes in replisome composition, we performed replication with fluorescently labelled DNA polymerases (LD655-gp5-trx). Under conditions without flow, we observe polymerase signal moving together with the lagging-strand product (Fig. 5A and fig. S12). To determine polymerase number, the mean fluorescence of single labelled polymerases was estimated prior to photobleaching (fig. S13). The half-life of labelled polymerases was calculated to be 10-fold longer than the observation time of replication demonstrating that photobleaching does not impact estimates of polymerase numbers (fig. S14). For the unconstrained DNA under no flow conditions, we estimate 3.75 ± 0.08 (mean ± SEM, n = 75 molecules) polymerases at the replisome consistent with previous estimates (Geertsema et al., 2014). Strikingly, for the constrained DNA, we observe a significant increase to 11.40 ± 0.22 (mean ± SEM, n = 56 molecules) polymerases at the replisome in the presence of the topological barrier (Table S1).

**Figure 5.**
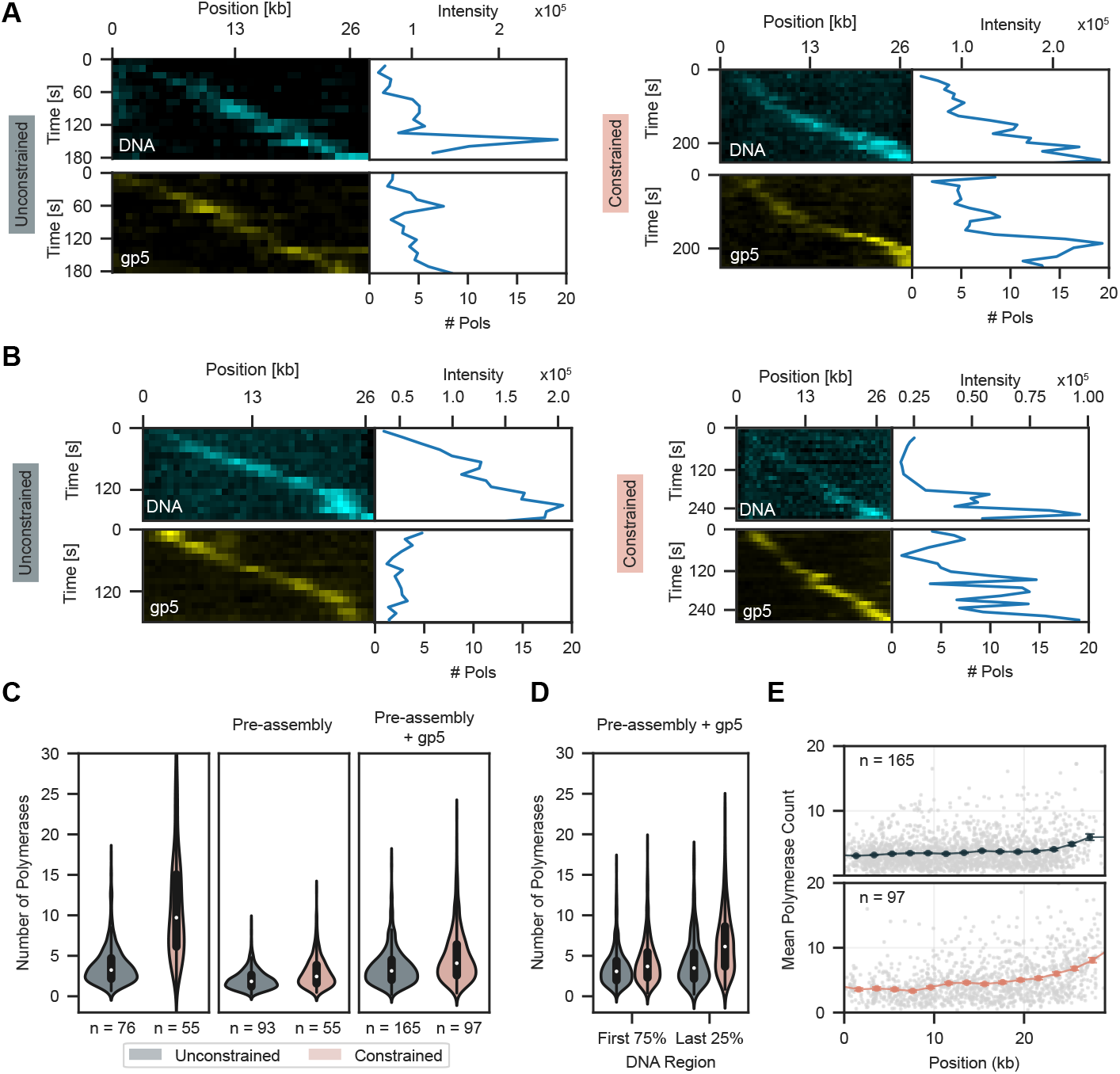
Topological strain results in polymerase accumulation. (**A**) Representative kymograph of replication of an unconstrained and constrained molecule in the linear configuration. (**B**) Representative kymograph of replication on an unconstrained and constrained molecule in linear configuration under pre-assembly condition with additional polymerases. Top kymograph displays the lagging-strand product and bottom displays polymerases in panels A and B. (**C**) Comparison of polymerase number distribution at the replication fork for unconstrained and constrained molecules for excess protein, pre-assembly, and pre-assembly with additional gp5. (**D**) Violin plot of polymerase numbers for unconstrained and constrained molecules under pre-assembly condition, including additional gp5, depending on position split in the first 75% and last 25% of replication substrate. (**E**) Sliding window analysis of number of polymerases for unconstrained (top) and constrained (bottom), depending on the replisome position on DNA. Grey data points represent the single measurements. Solid points represent the window means, with SEM indicated by the error bars.

The significant increase in polymerases beyond the six binding sites on gp4 (34)(Geertsema et al., 2014) prompted us to repeat our experiments using preassembled replisomes (fig. S15). With polymerases omitted after pre-assembly (fig. S16A), we observe 2.07 ± 0.04 (mean ± SEM) polymerases during replication of unconstrained molecules and 2.92 ± 0.08 (mean ± SEM) polymerases during replication of constrained molecules. Notably, omission of polymerases leads to a reduction in processivity for unconstrained molecules to 44 kb (95% confidence interval from 39 to 52, n = 178 molecules) and a further reduction for constrained molecules to 36 kb (95% confidence interval from 29 to 47, n = 76 molecules) (fig. S16).

Next, we performed experiments with pre-assembled replisomes with gp5 and gp2.5 included in solution during replication (fig. S17). Under these conditions, we observed 3.61 ± 0.05 (mean ± SEM) polymerases for the unconstrained molecules and 4.91 ± 0.09 (mean ± SEM) polymerases for the constrained molecules. The processivity of the unconstrained molecules increased modestly to 52 kb (95% confidence interval from 46 to 61, n = 194 molecules). In contrast, the processivity for constrained molecules remained unchanged when compared with the condition where polymerases were omitted.

The increase in polymerases we observe for the constrained molecules across all conditions (Fig. 5C) suggests excessive fork rotation disrupts polymerase-helicase coupling and leads to the recruitment of additional polymerases. Over time, this results in the accumulation of polymerases at the replication fork (Fig. 5D, E). These observations are consistent with a recent report that excess polymerases in solution help support continued replication under torsion (Jia et al., 2024a). The observed number of polymerases under pre-assembly conditions are below the six binding sites on gp4, suggesting the additional polymerases are associated with the replisome. In contrast, the number of polymerases during replication of constrained molecules in the absence of pre-assembly was beyond the six binding sites available on gp4. This could result from the loading of a second gp4 behind the active replisome that could recruit additional polymerases. Alternatively, lagging-strand polymerases released from the replisome may remain associated for long time periods after completion of synthesis as has been previously reported (Duderstadt et al., 2016). Since the lagging-strand product resides as a blob at the same location as the replisome, there released polymerases would be included in our estimates.

## Discussion

Our observations demonstrate that a rotational mode of replisome operation provides an intrinsic pathway to overcome topological challenges but results in negative consequences for replisome coordination. It remains to be resolved whether replisome rotation serves only as a fail-safe pathway during periods of acute stress or is a mechanical feature of normal replication. Topological barriers frequently form on regions of chromosomes where accumulation of overwinding outpaces topoisomerase activity. This occurs in highly transcribed regions that are rapidly overwound ahead of advancing RNA polymerases(Lang and Merrikh, 2021; Garcia-Muse and Aguilera, 2016; Janissen et al., 2024). During head-on encounters with the replisome the rate of overwinding is further increased and binding sites available for topoisomerases are reduced (Lang and Merrikh, 2021; Lang et al., 2017). Similar challenges occur at topological boundaries and near chromosome ends (Guo et al., 2021; Ullsperger et al., 1995; Stracy et al., 2019; Postow et al., 1999; Keszthelyi et al., 2016). Many common forms of stress can further complicate these issues (Branzei and Foiani, 2010). Therefore, topological barriers are clearly a major frequent challenge to DNA replication. Interestingly, two recent studies using both optical trapping and angular optical trapping have demonstrated that DNA torsion is a dynamic regulator of DNA replication stalling and reactivation (Jia et al., 2024a,b). However, live-cell imaging in *E. coli* has revealed that while topoisomerases accumulate at the replication fork, their number remains insufficient to keep up with the overwinding created by the replisome (Stracy et al., 2019). This latter observation opens up the possibility that replisomes may not only rotate during encounters with acute topological barriers but also during normal operation.

Dynamic exchange of polymerases provides robustness during DNA replication with the frequency adapting in response to environmental challenges (Loparo et al., 2011; Lewis et al., 2017, 2020; Tuan et al., 2022; Li et al., 2019; Kapadia et al., 2020; Beattie et al., 2017; Liao et al., 2016). Changes in polymerase coupling dynamics have been hypothesized to serve as a mechanical response pathway during encounters with torsional strain (Ullsperger et al., 1995). Normally, lagging-strand synthesis would be expected to continue until encountering the previously synthesized Okazaki fragment or a signal triggers early release (Li and Marians, 2000; Yang et al., 2006). In this scenario, the lagging-strand polymerase would dissociated from its gp4 binding site and continue Okazaki fragment synthesis behind the replisome. This type of premature release pathway was previously proposed as a mechanism to relieve torsional strain during rolling-circle replication (Kurth et al., 2013).

We observe that placement of a topological barrier directly ahead of the replication fork results in polymerase accumulation. This observation is consistent with torsional strain leading to more frequent polymerase dissociation from either the helicase or the template. Previous experimental observations and computational modeling have revealed that dissociation events from either site can facilitate the recruitment of additional polymerases (Geertsema et al., 2014; Åberg et al., 2016; Geertsema and van Oijen, 2013). Interestingly, modeling predicts that polymerases rapidly rebind under preassembly conditions, whereas exchange pathways are activated by polymerases in solution. Our observations are consistent with these predictions. However, further studies will be required to determine the occupancy of specific polymerase binding sites at the replication fork over time.

Eukaryotes have evolved accessory factors, such as the fork protection complex, that provide an early warning system to regulate replisome speed and reduce excessive fork rotation that can lead to uncoupling of daughter-strand synthesis and replication fork collapse (Keszthelyi et al., 2016; Schalbetter et al., 2015). Interestingly, chromatin can also help reduce the topological stress on the replication fork by acting as a shock absorber for moderate increases in overwinding (Le et al., 2019). These features could provide eukaryotes with more time for topoisomerases and the replisome to react and ensure fork rotation is limited in duration and rate to reduce the risk of disruption to replisome coordination. Nevertheless, there are many regions on chromosomes where the rate of overwinding can exceed what can be absorbed by chromatin. Stalling and replication fork collapse have been reported under these scenarios (Keszthelyi et al., 2016; Schalbetter et al., 2015). This highlights the importance of understanding intrinsic pathways that allow replisomes to cope with topological challenges. Further studies conducted with reconstituted eukaryotic replisomes are needed to determine exactly how nucleosomes modulate fork rotation dynamics.

Transverse flow imaging offers new opportunities to discover the dynamics at the replication fork that ensure robust genome duplication. By spatially resolving the parental, leading, and lagging strands in real-time during DNA replication, the approach opens up the possibility of time-resolved, strand-specific tracking of not only replication factors but also a wide array of other machineries that must coordinate their activities with the replisome. These events could be further evaluated in the context of diverse strand-specific obstacles frequently encountered on chromosomes. The approach can be easily adapted to studies of eukaryotic replication through small modifications to existing experimental platforms (Lewis et al., 2020). Taken together, this observation potential provides a powerful tool to discover the dynamics that support robust chromosome replication throughout the domains of life.

## Materials and Methods

### Gp4 purification

Recombinant gp4 proteins were overexpressed and purified using established protocols with necessary modifications (Jaisser et al., 1993; Notarnicola et al., 1995; Lee and Richardson, 2001). *E. coli* strain HMS 174(DE3), harboring a plasmid encoding the gp4 protein, was grown in LB medium to an optical density (OD600) of 1.0. Protein expression was induced by adding IPTG to a final concentration of 1 mM, and cultures were incubated for an additional 3 hours at 37 °C. Cells were harvested by centrifugation and resuspended in lysis buffer (20 mM Tris-HCl, pH 7.5, 5 mM EDTA, 0.1 M NaCl, 1 mM phenylmethylsulfonyl fluoride). Cell lysis was performed using a cell disruptor with a pressure of 25,000 psi. The lysate was clarified by centrifugation at 22,404 × g for 30 minutes, and the supernatant was treated with polyethylene glycol (PEG 4000) to a final concentration of 10%. The resulting precipitate was collected by centrifugation at 6,000 × g for 20 minutes, resuspended in binding buffer (20 mM potassium phosphate, pH 6.8, 1 mM EDTA, 1 mM DTT, 10% glycerol), and subjected to phosphocellulose chromatography. Bound proteins were eluted with a gradient of KCl (0.02–1 M), and fractions containing gp4 protein were pooled based on SDS-PAGE analysis. To further purify the protein, MgCl_2_ was added to the pooled fractions to a final concentration of 10 mM, and the solution was loaded onto an ATP-agarose affinity column. Protein was eluted using a buffer containing 20 mM potassium phosphate, pH 6.8, 20 mM EDTA, 0.5 mM DTT, 10% glycerol, and 0.5 M KCl. The elution fractions from the ATP-agarose column were pooled, and the protein purity was confirmed by SDS-PAGE analysis. The concentration was adjusted to approximately 10 µM to ensure that the final concentration of the hexameric helicase after stock buffer dialysis was maintained above 15 µM, which is optimal for long-term storage. The final solutions were dialyzed against storage buffer (20 mM potassium phosphate, pH 7.5, 0.1 mM DTT, 0.1 mM EDTA, 50% glycerol) and stored at −20 °C until use.

### Gp2.5 purification

Recombinant gp2.5 proteins were overexpressed and purified using established protocols with necessary modifications (Wrede et al., 2002). *E. coli* BL21(DE3)pLysS cells harboring a plasmid encoding gp2.5 were grown in LB medium to an optical density (OD600) of 1.0. Cells were harvested by centrifugation and resuspended in lysis buffer (50 mM Tris-HCl, pH 7.5, 0.1 mM EDTA, 1 mM DTT, 10% glycerol, 1 M NaCl) prior to freezing. Before extraction, frozen cells were thawed on ice in a cold room overnight. Cells were lysed by incubation with 1 mg/mL lysozyme for 1 hour at 4°C with continuous mixing, followed by cell disruption using a cell disruptor at a pressure of 25,000 psi. Cellular debris was removed by centrifugation at 22,040 × g for 30 minutes. The clarified lysate was further ultracentrifuged at 176,672 × g for 40 minutes.

Polyethyleneimine (10% v/v stock solution, pH 7.5) was added to the supernatant to a final concentration of 0.1% v/v, and the solution was stirred at 4°C for 1 hour. The precipitated proteins were pelleted by centrifugation at 21,000 × g for 30 minutes. The pellet was resuspended in Buffer A (50 mM Tris-HCl, pH 7.5, 0.1 mM EDTA, 1 mM DTT, 10% glycerol) and centrifuged again at 21,000 × g for 30 minutes.

Ammonium sulfate was added to the resulting supernatant to reach a final concentration of 80% saturation, followed by stirring for 1 hour at 4°C. The precipitate was collected by centrifugation at 6,000 × g for 30 minutes, resuspended in Buffer A, and subjected to ultracentrifugation at 176,672 × g for 40 minutes. The cleared protein solution was filtered through a 0.45 µm filter. The filtrate was loaded onto an HQ Poros column equilibrated with Buffer A supplemented with 50 mM NaCl and eluted using a NaCl gradient (0.05–1 M) in Buffer B (Buffer A supplemented with 1 M NaCl). Fractions containing gp2.5 were identified by SDS-PAGE, pooled, and dialyzed against dialysis buffer (50 mM Tris-HCl, pH 7.5, 0.1 mM EDTA, 1 mM DTT, 50% glycerol). The dialyzed proteins were stored at 80°C until use.

### LD655-gp5/trx purification and labeling

The expression and purification of T7 bacteriophage gp5/trx was based on the previously described protocol by Johnson and Richardson (Johnson and Richardson, 2003). Internally YBBR-labeled gp5 was cloned into a pRSFDuet vector. The YBBR tag (DSLEFIASKLA) was introduced between I464 and T465. Thioredoxin (trx) was cloned into a pET Duet vector. The plasmids were co-transformed into *E. coli* BL21 star cells. Cells were grown in 2 liters of TB medium containing Amp, Kan, and K salts at 37°C until reaching an OD600 of 1.0. Protein production was induced with 1 mM IPTG and incubated at 37°C for an additional 4 hours. All subsequent purification steps were performed on ice or at 4°C. The cells were harvested by centrifugation (4000 x g, 15 min), washed with PBS, and centrifuged again (4000 x g, 15 min). The resulting cell pellet was frozen in liquid nitrogen. To support cell lysis, the frozen cell pellet underwent a freeze-thaw cycle by thawing in a water bath, refreezing in liquid nitrogen, and thawing again. The pellet was resuspended in lysis buffer (25 mM HEPES-KOH, pH 8.0, 1 mM DTT, 5% (v/v) glycerol, 500 mM KCl, 20 mM imidazole). The lysis mix was supplemented with 1x protease inhibitor cocktail, 1x lysozyme, and 1x DNAase I. The cells were lysed by three rounds of sonication (5 minutes, 4 cycles, 30% power). The cell lysate was cleared by centrifugation at (27,000 x g, 30 min).

The cell lysate was applied to a HisTrap (5 ml) column equilibrated in lysis buffer. After sample application, the column was washed sequentially with lysis buffer and wash/desalting buffer (25 mM HEPES-KOH, pH 8.0, 1 mM DTT, 5% (v/v) glycerol, 200 mM KCl). The protein was then eluted using a gradient from 20 mM to 250 mM imidazole (Elution buffer: 25 mM HEPES-KOH, pH 8.0, 1 mM DTT, 5% (v/v) glycerol, 200 mM KCl, 250 mM imidazole). Peak fractions were pooled based on SDS-PAGE analysis and concentrated using a MWCO 10,000 Amicon Ultra Centrifugal Filter unit. The concentrated sample was applied to a HiPrep 26/10 Desalting column equilibrated in wash/desalting buffer to remove imidazole. The fractions were pooled based on SDSPAGE analysis and incubated overnight at 4°C with TEV protease at a 1:50 ratio and 1 mg DNase to cleave the His-tag. The sample was subsequently applied to a His-Trap (5 ml) column equilibrated in wash/desalting buffer, and the flow-through was collected. Next, the sample was loaded onto a HiTrap Heparin (5 ml) column equilibrated in wash/desalting buffer. The protein was eluted using a salt gradient from 200 mM to 1 M KCl. Peak fractions were pooled and concentrated using a MWCO 10,000 Amicon Ultra Centrifugal Filter unit. The sample was then applied to a Superdex 200 Increase 10/300 gel filtration column equilibrated in gel filtration (GF) buffer (25 mM HEPES-KOH, pH 8.0, 1 mM DTT, 10% (v/v) glycerol, 150 mM KCl, 0.1 mM EDTA). The protein eluted as a single symmetric peak, corresponding to an approximate molecular weight of 95 kDa (a 1:1 complex of gp5 and trx). Peak fractions were pooled based on SDS-PAGE analysis and spin concentrated.

To produce LD655-gp5 in complex with Trx, YBBRgp5/Trx was mixed with SFP synthase and LD655-CoA (lumidyne) at a molar ratio of 1:1:1.5 in GF buffer supplemented with 10 mM MgCl_2_ and incubated at 30°C for 2 hours. The sample was then applied to a Superdex 200 Increase 10/300 gel filtration column equilibrated in GF buffer. Peak fractions were pooled based on SDSPAGE analysis and concentrated using an Amicon Ultra Centrifugal Filter unit. Aliquots were snap-frozen and stored at −80°C. Labeling efficiency was estimated at approximately 94%, based on the extinction coefficients of the gp5/trx complex and LD655. The final protein concentration was determined using a spectrophotometer by measuring absorbance at 280 nm, while protein purity was assessed by the 260/280 absorbance ratio.

### DNA handle preparation with multiple biotins

For creating topologically constrained DNA substrates, handles with multiple biotins interacting with the slide surface were assembled. DNA sequence was amplified from lambda DNA via a PCR reaction. Each PCR reaction contained 50 ng of lambda DNA, 6 units of Phusion High-Fidelity DNA polymerase, PCR reaction buffer (1x HF Buffer), 200 µM of each dATP, dCTP, dGTP, and dTTP (from the dNTP bundle), 3% DMSO, and the appropriate forward and reverse primers (NotI: oligo1, oligo2; XhoI: oligo3, oligo4, table S2).

The PCR product was purified using the QIAGEN QIAquick PCR Purification Kit, following the manufacturer’s instructions. To incorporate biotin molecules into the DNA handle, the concentration of dTTP was reduced and partially replaced with Biotin-11-dUTP. Each PCR reaction contained 50 ng of the purified template from the previous reaction, 15 units of Taq DNA polymerase, 1x ThermoPol Reaction Buffer, 200 µM of dATP, dCTP, and dGTP, 130 µM dTTP, 70 µM Biotin-11dUTP, 3% DMSO, and the same forward and reverse primers (NotI: oligo1, oligo2; XhoI: oligo3, oligo4, table S2) as in the previous reaction.

The PCR product was purified using the QIAquick PCR Purification Kit, following the manufacturer’s instructions. The handles were digested with either 30 units of NotI-HF or XhoI in 1x rCut Smart Buffer containing 2 µg of DNA per reaction at 37°C for 3 hours, followed by heat inactivation at 65°C for 20 minutes to stop enzyme activity. The digested DNA was gel purified using gel electrophoresis (0.75% TBE agarose gel, 90 V, 1 hour). The handles were extracted using the QIAquick Gel Extraction Kit. Biotin-labeled handles were freshly prepared before each ligation reaction and kept on ice in a cold room until the ligation step.

### Preparation of linear, biotinylated DNA for single-molecule nicking frequency quantification

For single-molecule TIRF assays to quantify nicking frequency, a DNA substrate with topologically constrained ends was generated from pMSuperCos plasmid. Plasmids were isolated from *E. coli* DH5α using the QIAGEN Plasmid Maxi Kit following the manufacturer’s instructions. 125 µg of plasmid was digested overnight at 37°C with 125 units of XhoI and NotI-HF in 1x rCutSmart buffer. Before loading the sample onto a Sephacryl S1000 SF Tricorn 10/300 gel filtration column for size-exclusion chromatography, 0.1% SDS was added to stop the digestion reaction. The column was equilibrated with 10 mM Tris-HCl, pH 8.0, 300 mM NaCl, and 1 mM EDTA (S-1000 Buffer). Desired fractions containing the XhoI-NotI fragment were pooled, and precipitated with ethanol at −20°C. The linearized plasmid (XhoI-NotI fragment) was resuspended in 10 mM Tris-HCl, pH 8.0 and stored at 4°C until the ligation step.

To construct a torsionally constrained linear DNA substrate, handles containing multiple Biotin-11-dUTP were ligated onto the linearized DNA. The handles were prepared as described in the previous section. The reaction mixture contained 18 µg of linearized DNA, 2000 units of T4 DNA Ligase, 1x T4 DNA ligase Buffer, 2 mM ATP, 2 µg of NotI handle, and 2 µg of XhoI handle. The reaction was incubated overnight at 16°C. To remove excess handles from the final DNA product, the reaction mixture was loaded onto a Sephacryl S-1000 SF Tricorn 10/300 gel filtration column equilibrated with S1000 buffer. Peak fractions were pooled, precipitated with ethanol, and resuspended in 10 mM Tris-HCl, pH 8.0, and 0.1 mM EDTA. Aliquots of the final DNA substrate were snap-frozen in liquid nitrogen and stored at −80°C.

### Preparation of forked, biotinylated DNA for singlemolecule replication assays

For single-molecule TIRF assays, involving unconstrained and constrained substrates, a pre-primed DNA forked substrate was created to bind to the surface through streptavidin-biotin interactions. Depending on the number of biotin molecules present in a DNA substrate, the substrate was free to rotate around a single biotin interaction (unconstrained) or was fixed on the surface (constrained). Both substrates were created from the same 27 kb plasmid backbone in addition to different DNA modules. Similar approaches have been employed previously to investigate DNA replication (Lewis et al., 2020; Mueller et al., 2020) using fluorescence microscopy. A method for investigation of helicase unwinding on long DNA substrates containing one end with multiple digoxigenins has likewise been reported (McClymont Cameron, 2024).

Plasmids (pMSuperCos) were isolated from *E. coli* DH5α using the QIAGEN Plasmid Maxi Kit. The purified plasmid was then digested with XbaI and XhoI. The digestion reaction, which contained 125 µg of plasmid DNA, 125 units each of XhoI and XbaI, and 1x rCutSmart buffer, was performed at 37°C overnight. The reaction was supplemented with 0.1% SDS to stop the digest reaction. XhoI-XbaI fragments were purified by gel filtration using a Sephacryl S-1000 SF Tricorn 10/300 gel filtration column equilibrated with S-1000 buffer. Ethanol precipitation was then performed, and DNA pellet was resuspended in Tris-HCl, pH 8.0, and stored on ice until the ligation.

Pre-primed DNA fork containing XbaI matching sequence was created by annealing oligos (oligo7, oligo8, oligo9, Table S2) in a ratio of 1:6:60 in Duplex Buffer (30 mM HEPES-KCl, pH 7.5, 100 mM KOAc). The mixture was heated to 95°C for 5 min and then cooled to 4°C at 0.5°C/min. The end piece with a single biotin containing XhoI matching sequence was created by annealing oligos (oligo5, oligo6, Table S2) in a ratio of 6:1 and performing the same annealing procedure described for the fork. The biotin handle with multiple biotins matching XhoI was described in previous section (‘DNA handle preparation with multiple biotins’)

To create unconstrained and constrained substrates, a ligation reaction was performed containing 1x T4 DNA Ligase Buffer with 2000 units of T4 DNA Ligase, 18 µg of XhoI-XbaI fragments and 2 mM ATP at 16°C overnight. To produce the unconstrained substrate, the ligation reaction additionally included the end piece with a single biotin (oligo5, oligo6, Table S2) and the preprimed fork (oligo7, oligo8, oligo9, S2) at 140 nM each. For the constrained substrate, the XhoI biotin handle (2 µg in total) and the single biotin fork (oligo7, oligo8, oligo9, S2, final concentration 140 nM) were additionally included in the reaction. The steps following the ligation reaction were identical for the following steps onward. Excess DNA handles were removed by a Sephacryl S-1000 SF Tricorn 10/300 gel filtration column equilibrated with the S-1000 buffer. The peak fractions containing the ligated substrate were pooled, precipitated with ethanol and reconstituted in Tris-HCl, pH 8.0, and 0.1 mM EDTA. Aliquots of the final DNA substrates were snap-frozen in liquid nitrogen and stored at −80°C.

### Single-molecule assays

#### PEG-Biotin microscope slides preparation

Glass coverslips (22 x 22 mm, Marienfeld) were first cleaned with a Zepto plasma cleaner, then transferred to a glass container filled with acetone containing 2% (v/v) 3-aminopropyltriethoxysilane and incubated for 1.5 minutes. The reaction was quenched by adding an excess of deionized water (ddH_2_O) and the coverslips were subsequently rinsed with (ddH_2_O), dried with compressed air, and baked at 110°C for 1 hour. For functionalization, coverslips were covered with a fresh solution of 0.6% (w/v) Biotin-PEG-Succinimidyl Carbonate (MW 5000) and 15% (w/v) mPEG-Succinimidyl Carbonate (MW 5000) in 0.1 M NaHCO_3_ and incubated overnight at room temperature. After incubation, the coverslips were rinsed with ddH_2_O, dried with compressed air and incubated again with a fresh Biotin-PEG/mPEG solution as described above. Functionalized PEF-Biotin microscope slides were stored under vacuum.

#### Flowcell preparation

A functionalized PEG-Biotin microscope slide was covered with 0.2 mg/ml streptavidin in blocking buffer (20 mM Tris-HCl, pH 7.5, 50 mM NaCl, 2 mM EDTA, 0.2 mg/ml BSA, 0.005% (v/v) Tween20) for 30 min. On top of the previously washed and dried slide, a polydimethylsiloxane (PDMS) block was placed to assemble a flow cell. The PDMS block had either the linear or the transverse flow cell configuration. The flow channels had a height of 0.1 mm and a width of 0.5 mm. Polyethylene tubes with an inner diameter of 0.58 mm were placed in the inlets and outlets of the PDMS. The flow channel was flushed with blocking buffer and incubated for 15 minutes.

The DNA tethering process varied depending on the flow cell configuration. For the linear flow cell configuration, 5 pM of forked, biotinylated DNA was introduced at a flow rate of 17 µl/min for 24 minutes in reaction buffer (Tris-HCl, pH 7.4, 50 mM Potassium Glutamate, 10 mM Magnesium chloride, 0.1 mg/ml BSA, 10 mM DTT) supplemented with 200 µM chloroquine. Unbound DNA was then washed out using reaction buffer supplemented with 300 µM ATP/CTP, 600 µM dNTPs, and 150 nM SYTOX Orange at a flow rate of 20 µl/min for 10 min. For the pre-assembly condition, the reaction buffer was supplemented with 300 µM ATP/CTP, 600 µM dATP, dTTP, dGTP, and 150 nM SYTOX Orange.

For the transverse flow cell configuration, inlet 2 and outlet 2 were clipped off with metal clips to prevent side flow. Then, 5 pM of forked, biotinylated DNA was introduced at a flow rate of 20 µl/min for 24 min in reaction buffer. Unbound DNA was washed out with reaction buffer containing 300 µM ATP/CTP, 600 µM dNTPs, and 150 nM SYTOX Orange at a flow rate of 20 µl/min for 10 min, while inlet 2 and outlet 2 remained clipped. The clips were then removed from inlet 2 and outlet 2, and inlet 1 and outlet 1 were clipped. Washing buffer was introduced at a flow rate of 20 µl/min for 5 min to ensure an even distribution of SYTOX Orange throughout the flow cell.

#### Single-molecule photoinduced nicking assay

To introduce negative supercoils in the linear, biotinylated DNA, the DNA application process in the transverse flow configuration was slightly modified. The workflow for inducing negative supercoils in DNA was adapted from the protocol by Ganji, Kim, van der Torre, Abbondanzieri and Dekker (Ganji et al., 2016). During the DNA tethering step, the linear, biotinylated DNA was mixed in reaction buffer containing 500 nM SYTOX Orange. The DNA was applied following the same procedure described for the transverse flow cell configuration, with the SYTOX Orange concentration reduced to 150 nM during the subsequent wash steps, which facilitated the formation of plectonemes. Different imaging conditions with varied laser powers were tested and compared.

#### Single-molecule replication assay in linear configuration

To initiate the T7 replication reaction in the linear configuration, a solution containing 2.5 nM gp4 (hexamer), 20 nM LD655-gp5/trx, and 750 nM gp2.5 in reaction buffer supplemented with 300 µM ATP/CTP, 600 µM dNTPs, and 150 nM SYTOX Orange was introduced into the prepared flow cell at a flow rate of 150 µl/min for 1.5 min. For experiments with DNA gyrase 5 nM was added to the T7 replication mix. The inlet and outlet tubes were then clipped to prevent any flow fluctuations or external force. The proteins were allowed to incubate for 0.5 minutes before starting image collection.

For the pre-assembly condition, a solution containing 2.5 nM gp4 (hexamer), 20 nM LD655-gp5/trx, and 750 nM gp2.5 in reaction buffer supplemented with 300 µM ATP/CTP, 600 µM dATP, dTTP, and dGTP, and 150 nM SYTOX Orange was introduced into the prepared flow cell at a flow rate of 150 µl/min for 1.5 min. The inlet and outlet tubes were then clipped, and the flow cell was incubated for 1.5 min. Depending on the pre-assembly condition, the flow cell was washed with replication buffer containing 300 µM ATP/CTP, 600 µM dNTPs, and 150 nM SYTOX Orange, or with replication buffer supplemented with an additional 20 nM gp5/trx and 750 nM gp2.5. The solution was flushed in at 150 µl/min. The inlet and outlet tubes were clipped, and data collection was started.

#### Imaging conditions

Single-molecule experiments were performed using a micromirror TIRF microscope from Mad City Labs (MCL, Madison, Wisconsin, USA) with custom modifications. The microscope was equipped with an Apo N TIRF 60 x oil-immersion TIRF objective (NA 1.49, Olympus). All experiments were performed in a temperature-controlled room at 22.5 ± 0.5 °C. SYTOX Orange and LD655 dyes were excited with a 532 nm and 637 nm laser (OBIS 532 nm LS 120 mW and OBIS 637 nm LX 100 mW, Coherent). An emission filter (ZET532/640 m, Chroma) was used to remove residual scattered light from excitation and to separate signals. Emission light was split at 635 nm (T635lpxr, Chroma) and collected on a Photometrics PrimeBSI sCMOS camera and later, an Andor iXon Life 888 EMCCD camera with comparable specifications. Protein or DNA was visualized sequentially every 5-10 s with a 100 ms integration time for 20 – 30 min. All microscope parts were controlled by Micromanager v2.0.0 (Edelstein et al., 2010; Schneider et al., 2012) and custom BeanShell scripts.

### Single-molecule data analysis

#### Image processing

All single-molecule raw data were processed in Fiji (Schindelin et al., 2012) using Molecule Archive Suite (Mars) commands (Huisjes et al., 2022). Stuck dots on the slide surface were used to correct for stage drift. For the linear configuration, a Single Molecule Archive was generated by tracking with subpixel resolution individual lagging strand products moving along the DNA. To generate a DNA Molecule Archive, individual DNA molecules were fit and checked for colocalization with individual molecule trajectories from the previous step. The DNA blob position on DNA versus time was fitted with a kinetic change-point algorithm (Hill et al., 2018) by assigning individual regions to distinguish between reaction and stalling.

#### Nick event analysis, quantification and fitting

To analyze nicking events, supercoiled DNA molecules were fitted using the ‘Object Tracker’ tool from Mars. Upon nick introduction, the area of the DNA molecule increases, which could be analyzed using the kinetic change-point step fitting. In addition, each molecule was inspected manually. Survival curves of the molecules were fitted using Python.

To create survival curves, the percentage of unnicked molecules at each time point was calculated by dividing the number of unnicked molecules at a specific time point by the total number of constrained molecules present at the start of the measurement. This percentage was then plotted against time. The half-life was determined by fitting the curves with an exponential decay model, where b represents the half-life of the curve and y0 denotes the initial quantity:

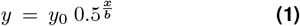

#### Spatial-temporal protein dynamics and kinetics in linear flow cell configuration

To analyze replication events in the linear flow cell configuration, the lagging strand product of active molecules was tracked with ‘Peak Tracker’, creating a Single Molecule Archive. Tracking was corrected for stage drift by identifying immobile peaks and using the coordinates as a reference. DNA molecules were fitted and tracking coordinates were transformed onto the reference frame of individual DNAs creating a DNA molecule Archive allowing the description of kinetics in terms of base pairs. An activity region was defined for each molecule and kinetics were fitted with the Kinetic change point. Molecules were sorted in subgroups. Molecules that exceeded the set timepoint for the 95% confidence interval were excluded. Rates lower than 15 bp/s (6-fold rate reduction compared to the mean rate) and durations >25s were considered pausing events. For the processivity, the endpoint of the final segment fitted by the kinetic change point was considered as the final position.

#### Processivity estimation using maximum likelihood estimation

Polymerase processivity was estimated using maximum likelihood estimation (MLE) for an exponential survival model with right-censoring. The method treats polymerase dissociation as a stochastic process where the probability of dissociation follows an exponential distribution characterized by a rate parameter λ. Observed dissociation events before the template end contribute to the likelihood through their probability density function, while polymerases completing the full template length are treated as right-censored observations contributing through their survival probability. The rate parameter λ is estimated by minimizing the negative loglikelihood function, and the mean processivity is calculated as 1/λ. Confidence intervals are derived using Fisher information, and the model’s goodness-of-fit is assessed by comparing predicted versus observed completion fractions at the template cutoff length.

#### DNA product shape analysis in the linear flow cell

A kymograph of each molecule was created using ‘DnaArchiveKymographBuilder’ tool from mars-kymograph by averaging 5 pixels on each side of the marked DNA molecule for every timestep. Pixel values were saved in a table. Results were smoothed with a sliding window (window size 2 pixels). The ‘threshold_otsu’ function from the scikit-image package was used to find the global threshold for each kymograph. Pixels below the threshold were set to 0 (black) while those above resulting in value 1 (white). Blob size of the DNA was the number of white pixels. The average size was calculated from 5 timepoints at the end of the reaction. The five timepoints were chosen from the interval between 10 and 5 timepoints preceding the final timepoint of the reaction.

#### Quantification of labelled gp5 protein and polymerase estimation

The average intensity of a labelled protein gp5 was quantified by analyzing bleaching steps in surface-immobilized molecules. The step size for each experiment was determined separately. Bleaching steps were fitted using kinetic change point step fitting. Images were beam-profile corrected. Protein spots were tracked using ‘Peak Tracker’ and simultaneously integrated (Inner radius 3 and outer radius 12) to measure protein signal. The resulting intensity value was divided by the estimated step size to calculate the number of polymerases.

#### Rate and processivity analysis for transverse flow

Transverse flow replication data were imported into Labkit (Arzt et al., 2022) and replication molecules were manually segmented. Strands of ‘Leading’, ‘Lagging’, and ‘Parental’ were traced in each frame. Segmentation files were imported into Mars using a custom importer (‘Transverse Flow Archive Builder’). Lengths of each segment are saved in pixels and converted to base pairs for each timepoint. After import into Mars, replication rates and processivities were determined in a manner similar to that used for the linear flow cell configuration. Kinetic changepoint analysis was applied to extract replication rates for the Leading, Lagging, and Parental strands. Manual segmentation of the individual strands was more challenging during fork rotation events because it was more difficult to accurately identify the location of the replisome without the three-way junction being visible. This can result in more variability in the rate distribution as displayed in Fig. 4D.

#### Analysis of lagging strand arm length dynamics

Lagging strand arm length over time was first tested using Dickey-Fuller unit root test (Perktold et al., 2024) to examine whether trace is non-stationary. Time traces were transformed into a stationary function by taking the difference between length at a timepoint t x(t) and length at previous timepoint x(t-1). Traces were tested with the augmented Dickey-Fuller unit root test to confirm stationarity. The variance of the length changes of the stationary function was calculated.

#### Estimating forces using the worm-like chain model

Force was estimated using the worm-like chain (WLC) model, where kB is the Boltzmann constant, T is the temperature (296.15 K), L is the contour length, A is the persistence length (50 nm) (Garcia et al., 2007; Bustamante et al., 1994), and z the current extension of the DNA for each time step.

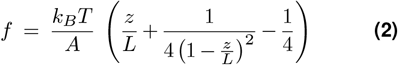

To estimate the force on the DNA arch consisting of leading and parental strand, contour length L was calculated by multiplying DNA length in bp times 0.34 nm (rise per basepair) and compared to the measured arch length. To estimate the force on the replisome, the extension of the lagging strand product was compared to the expected length based on the leading strand extension for same timepoint. Forces were grouped into bins of 3 kb, and a moving average with a step size of 1.5 kb was applied. Values below 5 kb were excluded from analysis. The median force for each bin was calculated and plotted. The resulting data were fitted with a line plot to illustrate the trend in force.

#### Quantification and statistical analysis

The number of observations (n) is indicated in the figure or figure legends. Errors in this study represent either the standard error of the mean (SEM) or the standard deviation (SD), as indicated. Python packages NumPy, pandas, matplotlib and seaborn were used for all statistical analysis.

## Acknowledgements

We are grateful to Margot Riggi of the Max Planck Institute of Biochemistry for help in making renderings of the replisome displayed in the manuscript. We would like to thank Fritz Simmel, Hannes Mustchler, Matthias Rief, and Martin Zacharias for critical feedback and support. TMR, LTR and KED were supported by a grant from the Deutsche Forschungsgemeinschaft (DFG, German Research Foundation) SFB863-11166240, a starting grant from the European Research Council (Project Number: 804098, REPLISOMEBYPASS), a consolidator grant from the European Research Council (Project Number: 101125005, ChromoMemInMotion), and the Max Planck Society.

## Author Contributions

Conceptualization, TMR, KED; Methodology, TMR, LTR, MT, KED; Investigation, TMR, LTR, KED; Visualization, TMR, KED; Funding acquisition, KED; Project administration, KED; Supervision, KED; Writing – original draft, TMR, MT, SMH, KED; Writing – review & editing, TMR, MT, SMH, KED

## Supplementary Information

**Figure S1.**
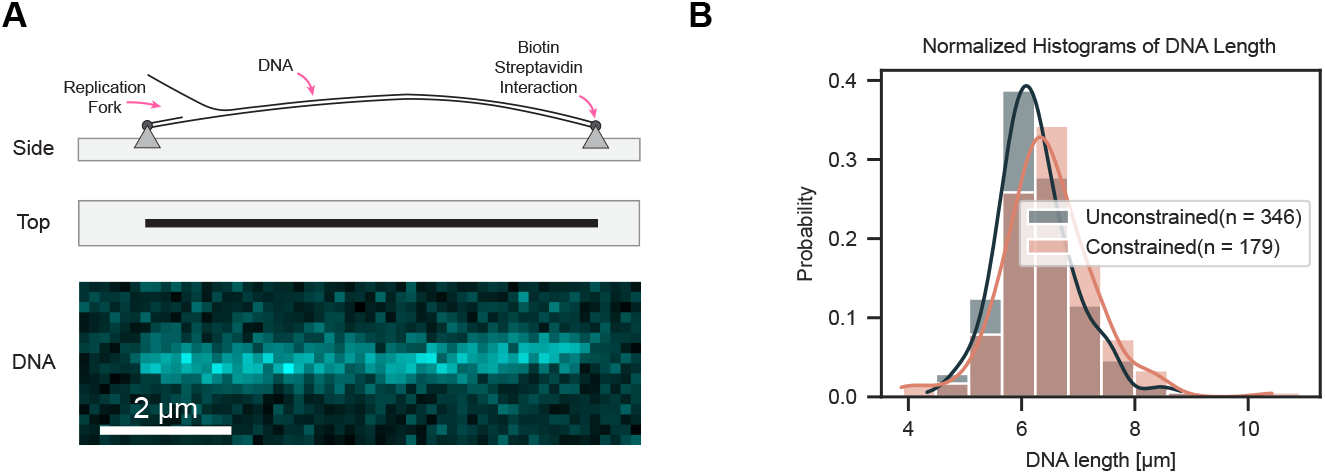
Molecule length distributions. (**A**) DNA attachment in linear flow cell configuration. Cartoon indicates replication fork where replication is initiated. DNA is tethered to the surface utilizing a biotin streptavidin interaction. (**B**) Length Distribution for the two substrate types in µm. Number of molecules is indicated in the legend. Unconstrained and constrained are stretched to 68±7% and 69±9% (Mean ± SD), respectively.

**Figure S2.**
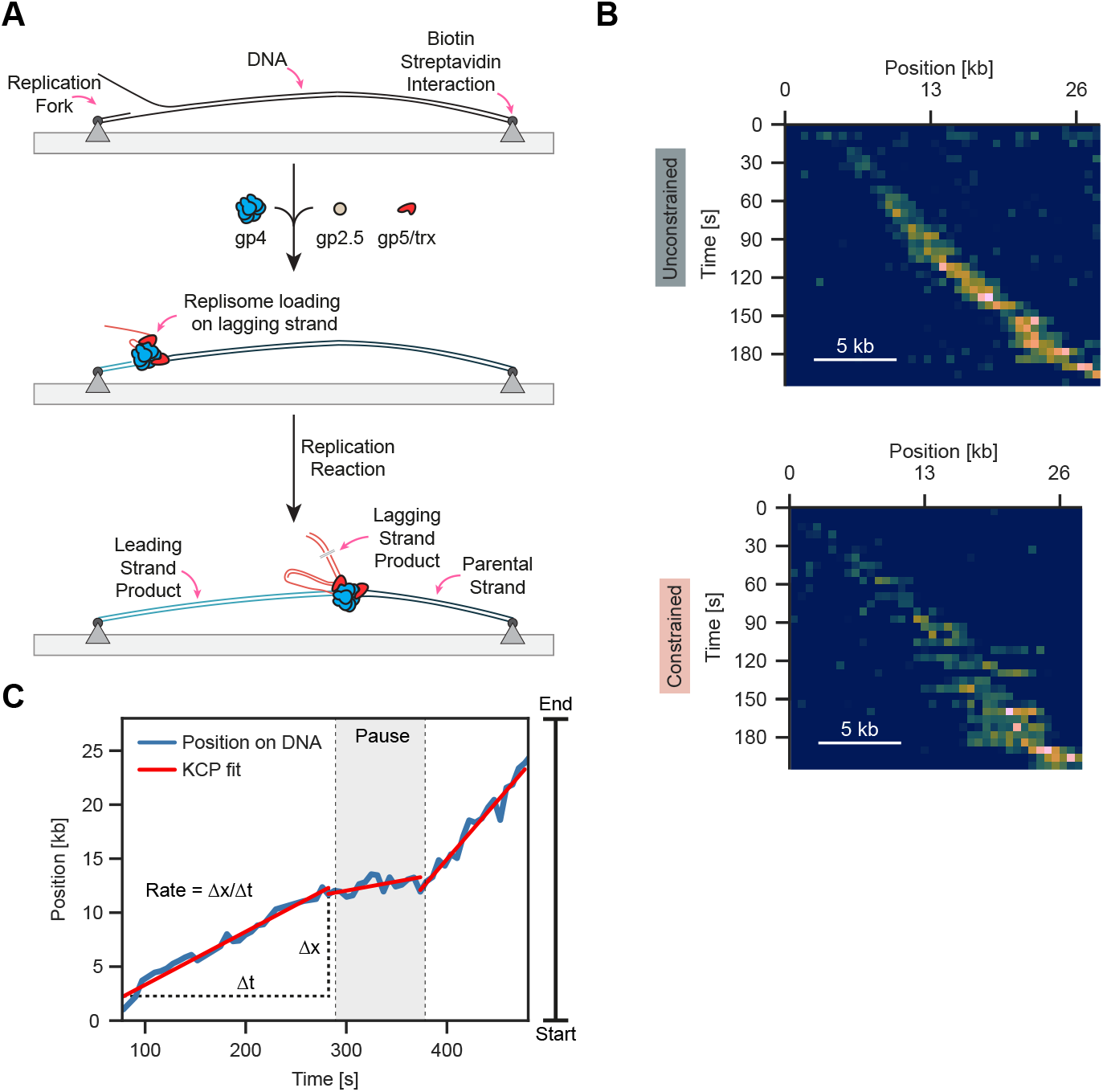
Replication using a linear flow configuration. (**A**) Schematic of replication. DNA is tethered to the surface using biotin-streptavidin interactions. T7 replication components initiate replication on the preformed fork. Leading strand product forms behind the replisome while the lagging strand product forms a blob at the location of the replisome used to track progression. (**B**) Example kymographs for single-molecule replication on unconstrained and constrained molecules. (**C**) Single-molecule tracking of lagging-strand product resulting in a time trace containing replication rates and processivities.

**Figure S3.**
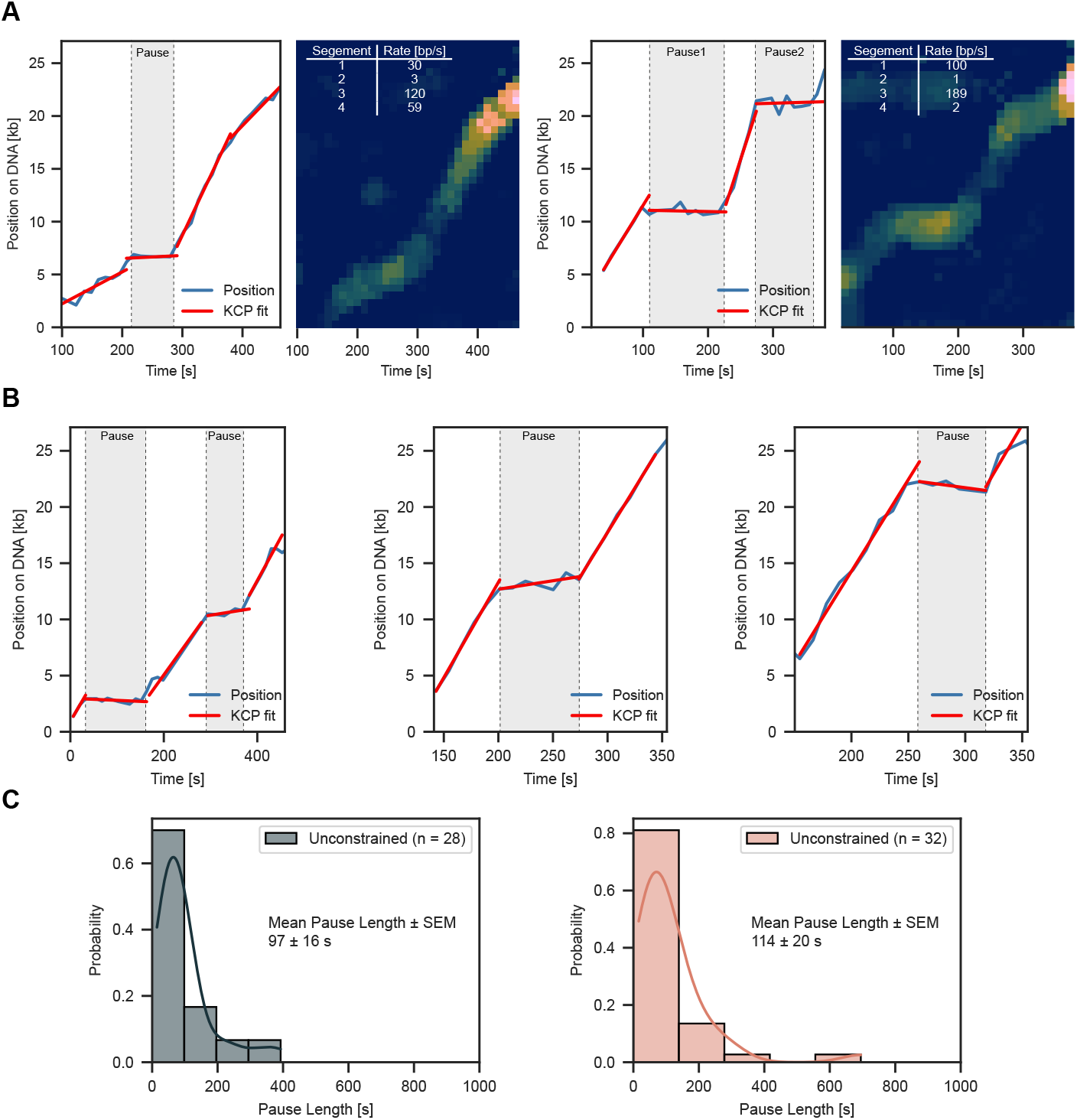
Pausing behavior. (**A**) Example traces and kymographs showing single or double pausing events during replication. The table represents rates fitted using kinetic change point analysis. (**B**) Additional example traces for pausing. (**C**) Representative pause duration distribution for unconstrained and constrained molecules.

**Figure S4.**
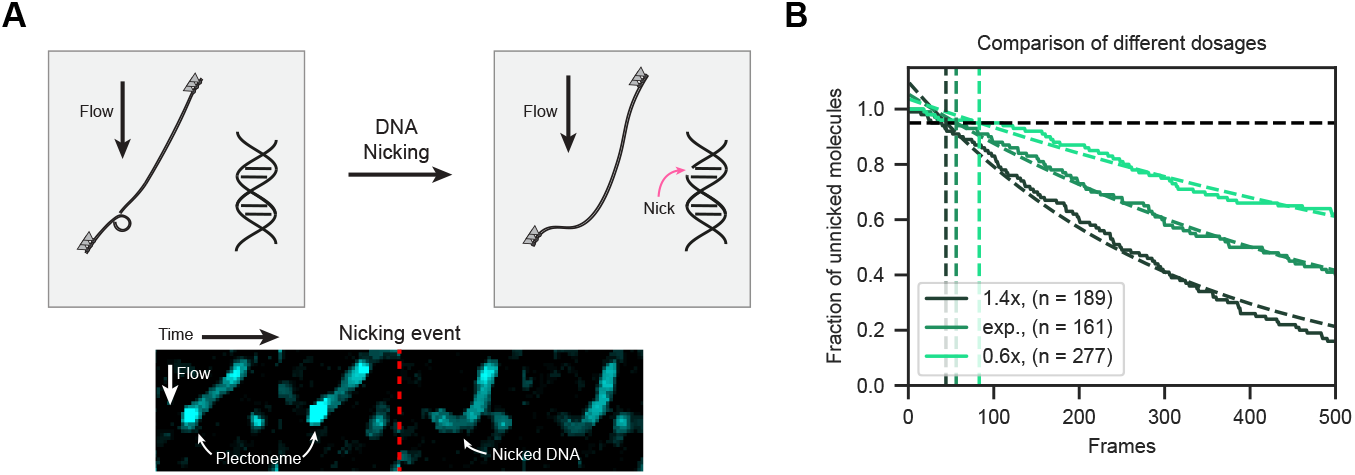
Nick detection assay for quantification of laser-induced damage. (**A**) Concept of nicking assay to control for imaging condition introducing nicks in individual strands. DNA is supercoiled and upon nick introduction supercoils are relaxed and result in an increase in the overall size of the DNA shape. (**B**) Survival curve for unnicked molecules over time fit with an exponential decay curve. Three different laser settings were compared. 1.4 times and 0.6 times the laser setting of the experimental condition (‘exp’) are displayed.

**Figure S5.**
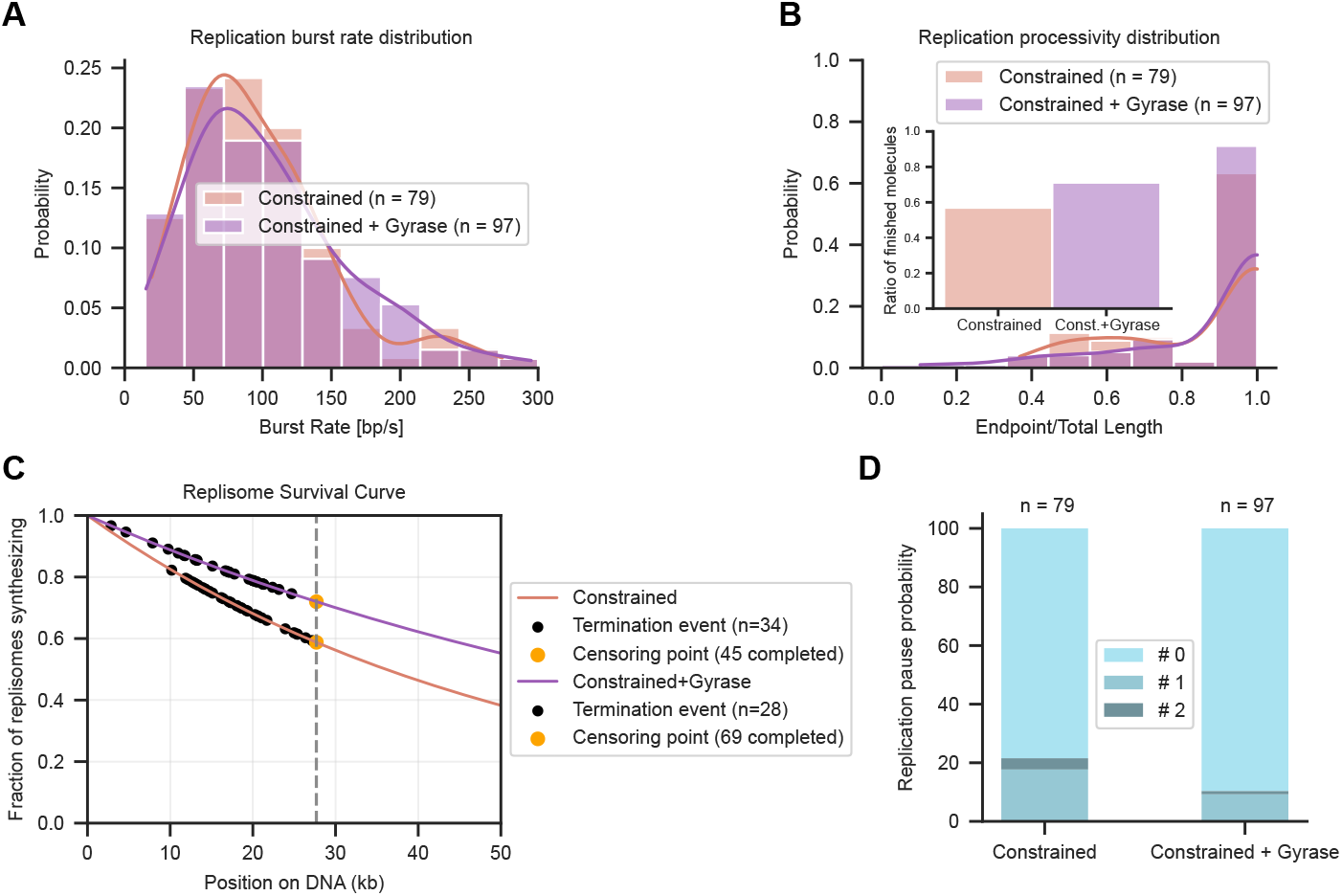
Visualizing replisome progression under topological strain with DNA gyrase. (**A**) Replication burst rate distribution for constrained and constrained including DNA gyrase. (**B**) Replication processivity distribution for constrained and constrained including DNA gyrase. Inset displays the fraction of molecules that replicated to the end. (**C**) Replication processivity estimation using maximum likelihood estimation. (**D**) Pause probabilities for constrained and constrained including DNA gyrase.

**Figure S6.**
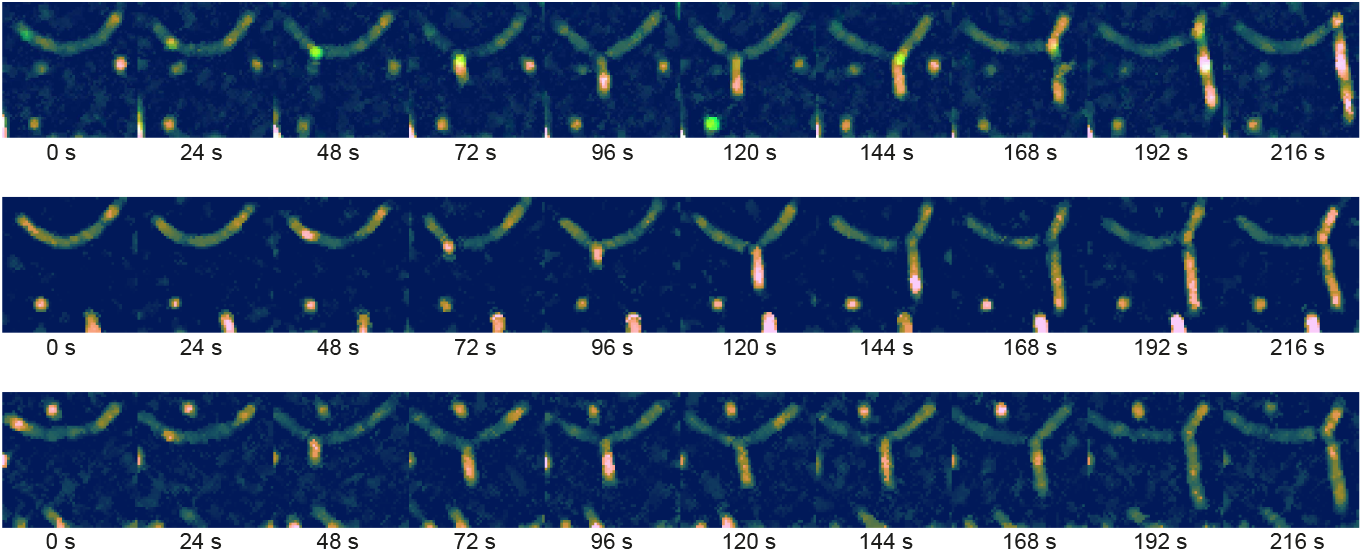
Representative molecules for transverse flow imaging of DNA replication (unconstrained). Three representative montages for replication reactions with transverse flow for unconstrained molecules.

**Figure S7.**
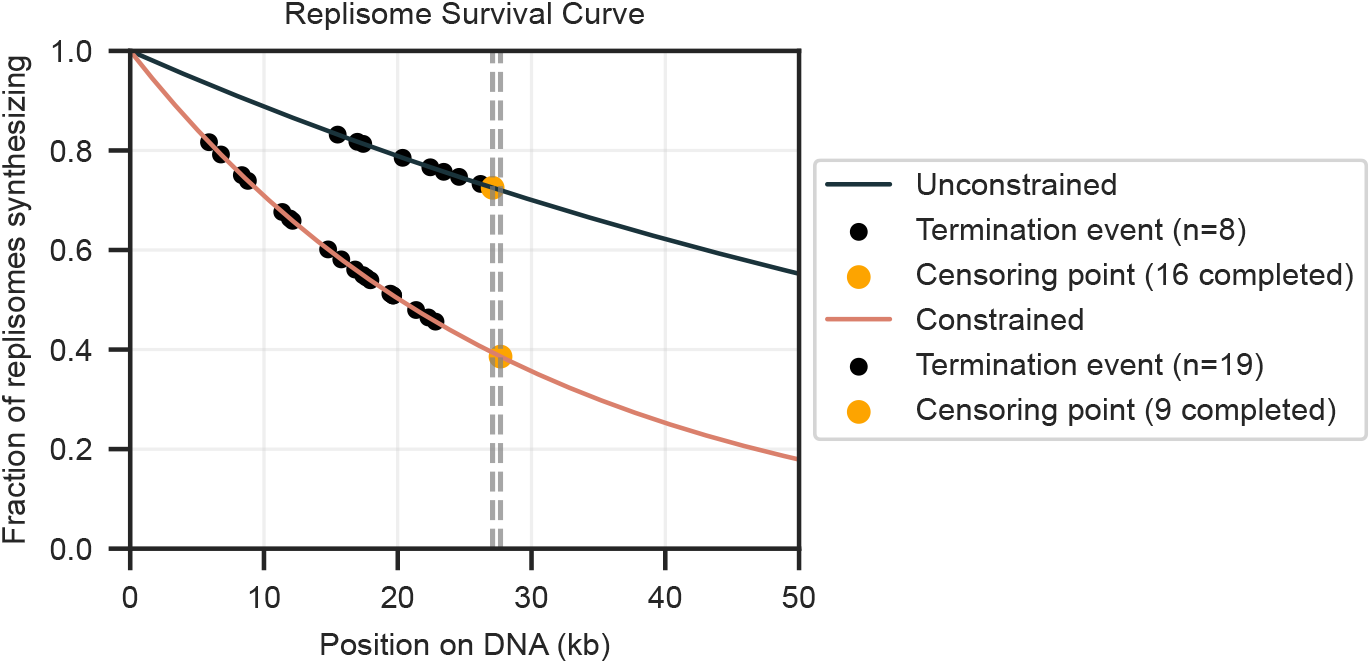
Processivity for transverse flow. Replication processivity using maximum likelihood estimation.

**Figure S8.**
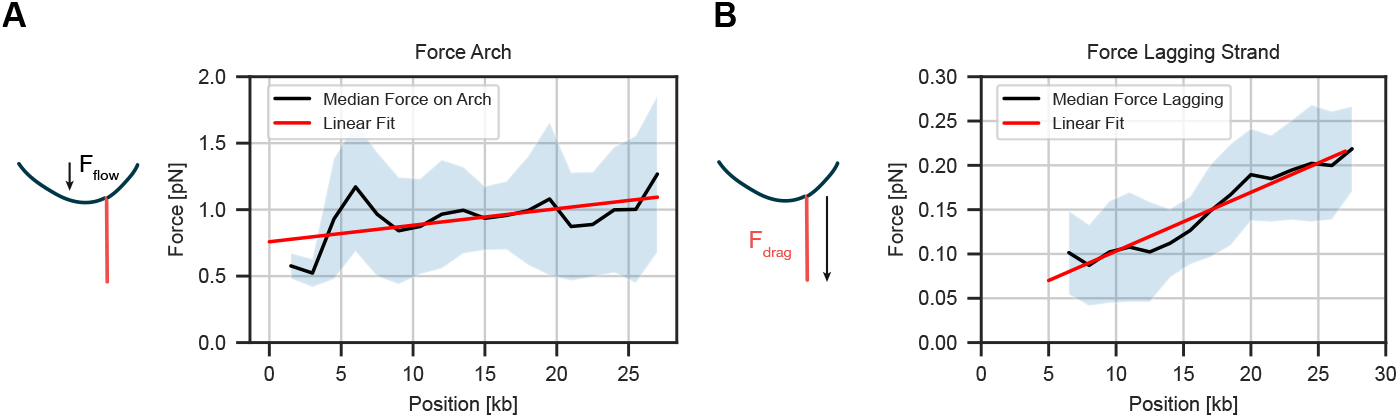
Estimation of applied forces from transverse flow. (**A**) Force on the arch applied by the flow. Force was calculated comparing arch extension to the contour length of the DNA substrate. (**B**) Applied force on the lagging-strand product due to transverse flow as a function of product position. Worm-like chain model was used for force estimation and a linear fit was added. Standard error of the mean indicates error margin.

**Figure S9.**
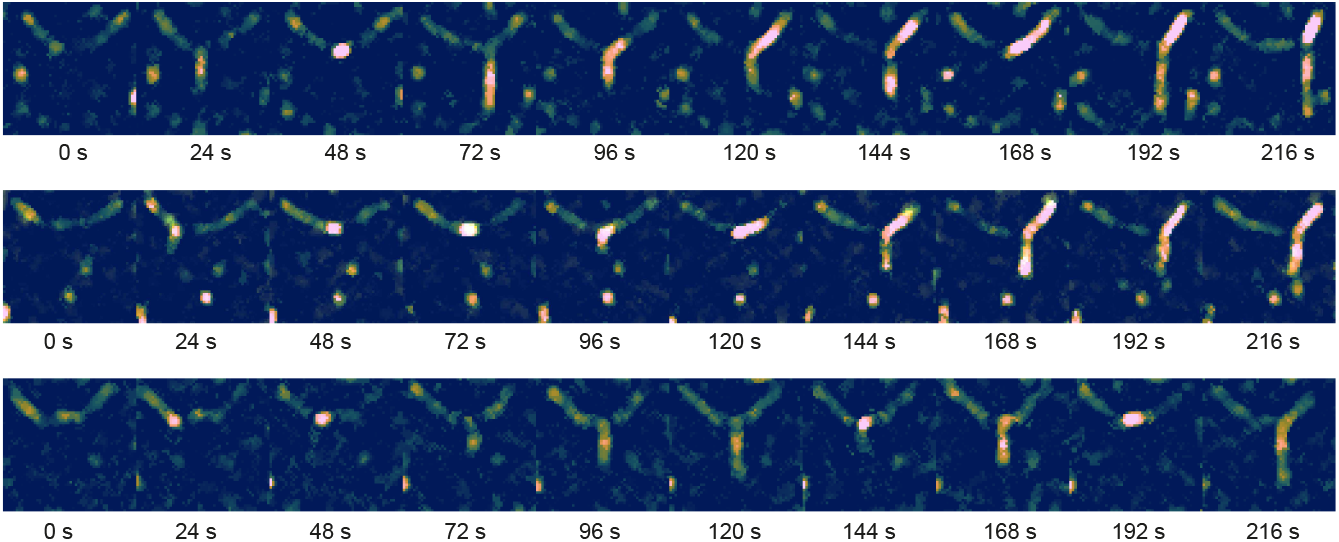
Representative molecules for transverse flow imaging of DNA replication (constrained). Three representative montages for replication reactions with transverse flow for constrained molecules.

**Figure S10.**
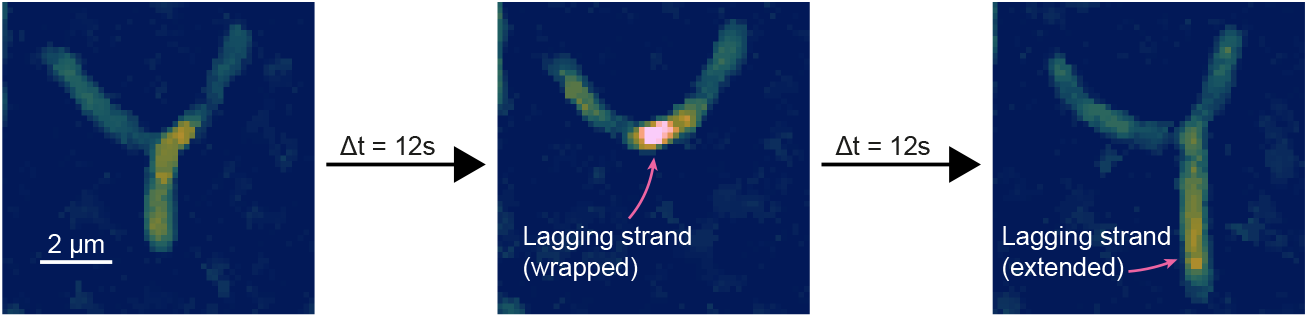
Montage of time points showing fork rotation. Three consecutive time points first showing an elongated lagging strand product, followed by complete wrapping (DNA blob) and finally re-extension of the lagging-strand replication product.

**Figure S11.**
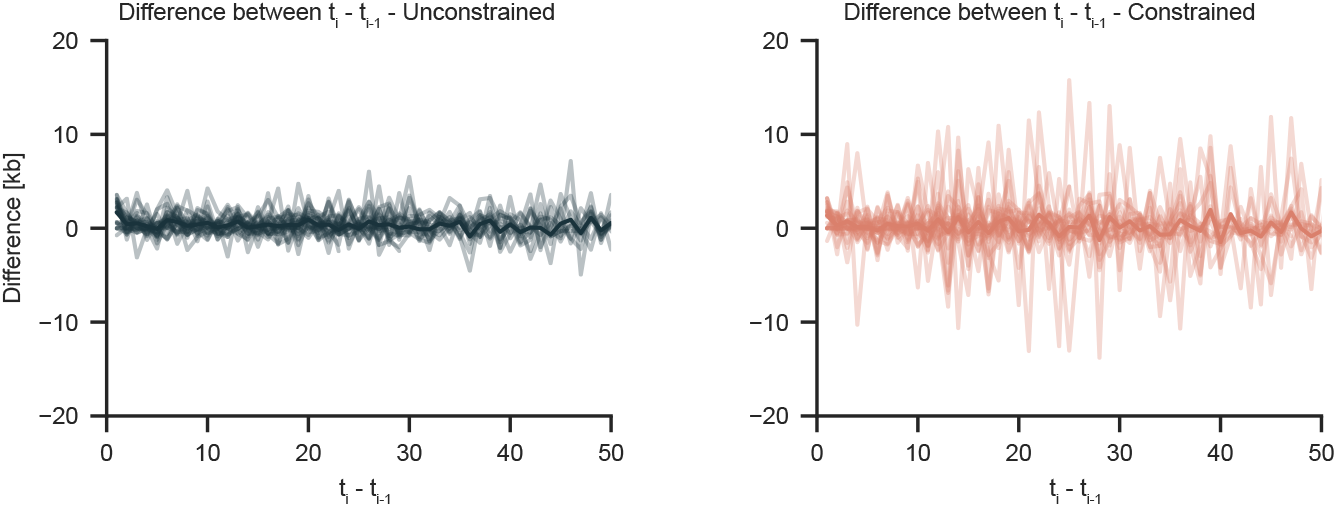
Transformed lagging strand length. Transformation of lagging strand length over time (Fig. 4F) from non-stationary function to stationary function. To transform, the difference is taken for each point by subtracting the value from the previous timepoint.

**Figure S12.**
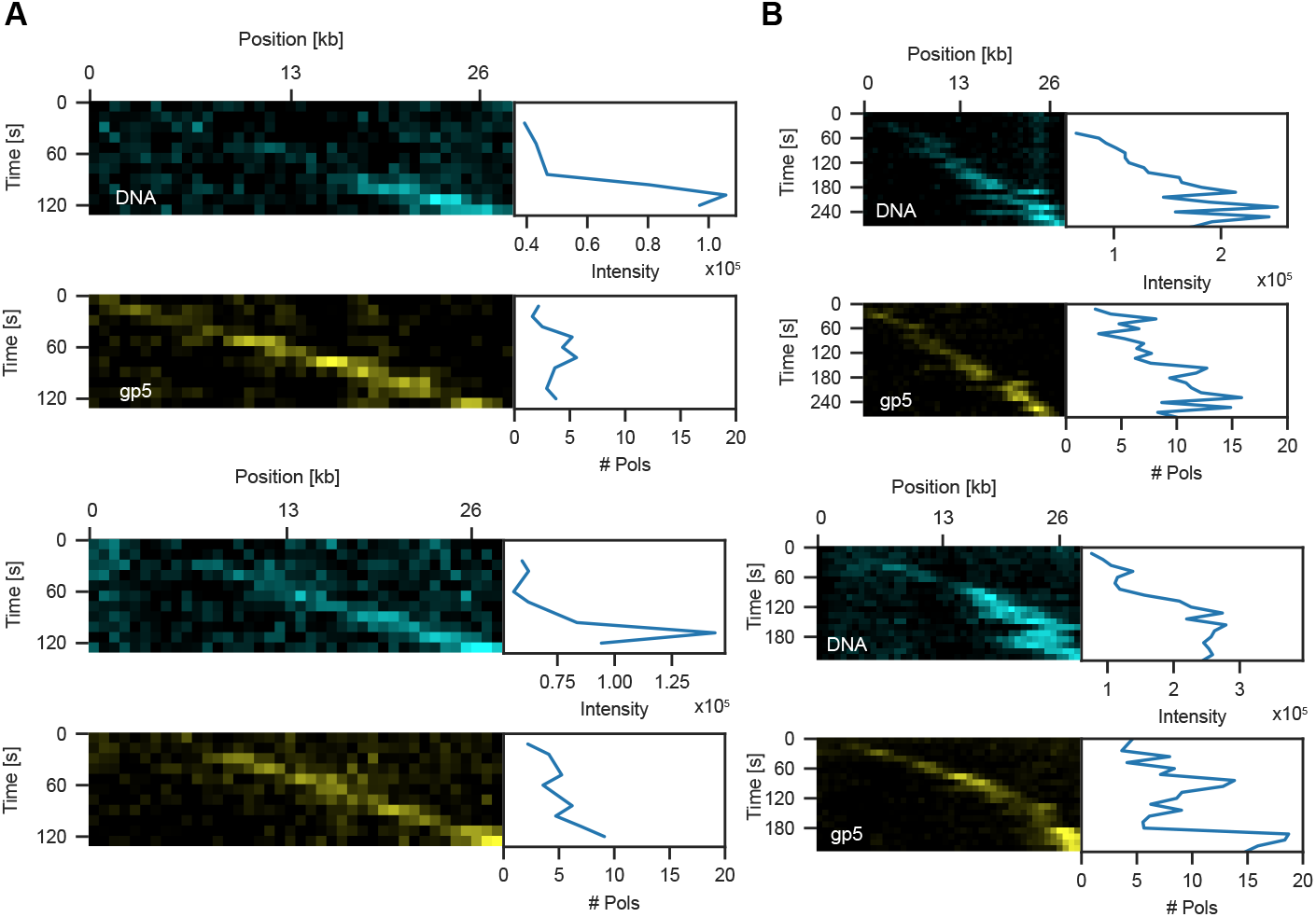
Representative molecules for labeled polymerases during DNA replication. (**A**) DNA replication events in the absence of flow on unconstrained molecules. (**B**) DNA replication events in the absence of flow on constrained molecules. Stained DNA is displayed in cyan on the top and polymerase signal is displayed in yellow on the bottom.

**Figure S13.**
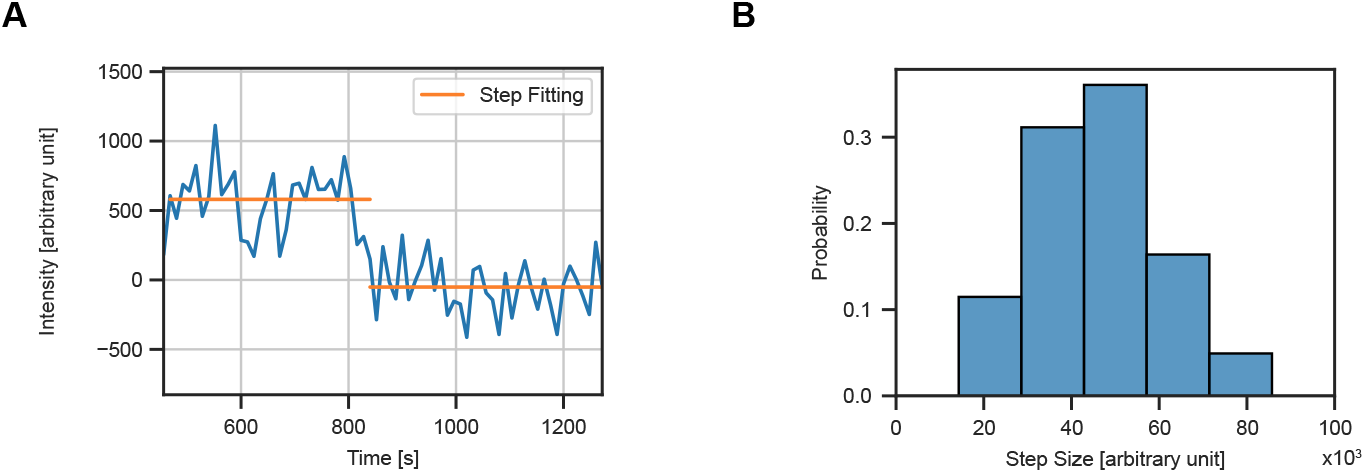
Estimation of fluorescent signal of a single labeled polymerase. (**A**) Bleach step of a surface immobilized polymerase. Step was fitted using change point analysis. (**B**) Distribution of different step sizes for one single molecule dataset.

**Figure S14.**
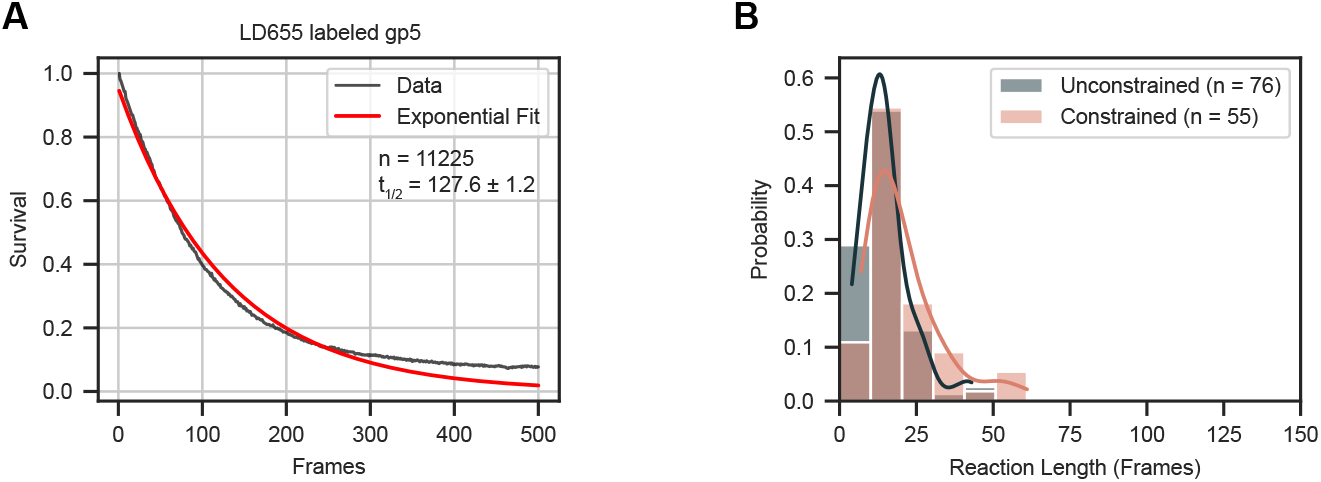
Lifetime of single LD655 labeled gp5. (**A**) Lifetime of single, surface bound LD655 labelled gp5 dyes. These were used to determine the number of frames until half of the dyes bleached. Half-life shown as mean ± SD. (**B**) Distribution of reaction times for replication events for unconstrained (15 ± 1 frames, mean ± SEM) and constrained (21 ± 2 frames, mean ± SEM).

**Figure S15.**
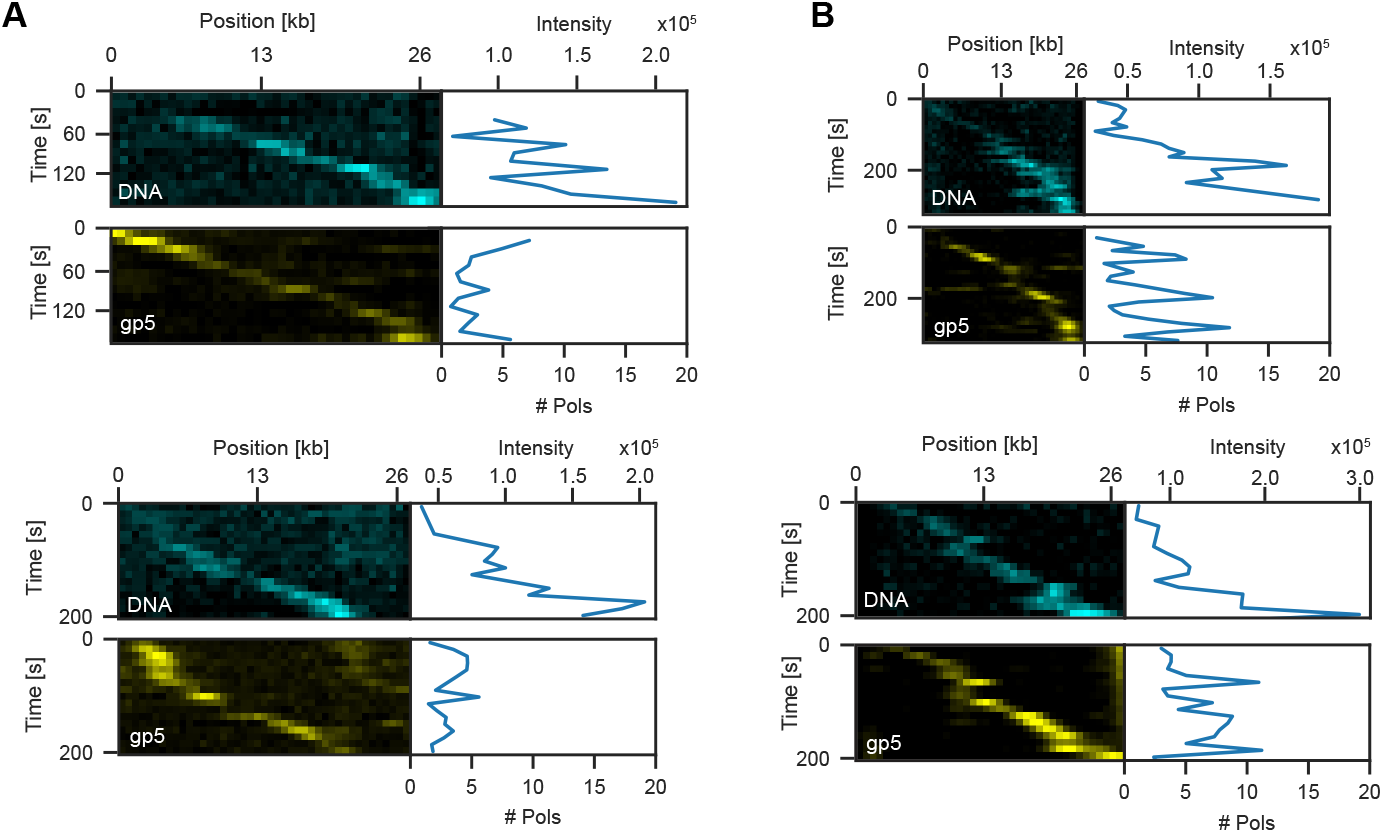
Representative molecules for labelled polymerases during DNA replication under pre-assembly conditions with additional gp5. (**A**) DNA replication events in the absence of flow on unconstrained molecules. (**B**) DNA replication events in the absence of flow on constrained molecules. Stained DNA is displayed in cyan on the top and polymerase signal is displayed in yellow on the bottom.

**Figure S16.**
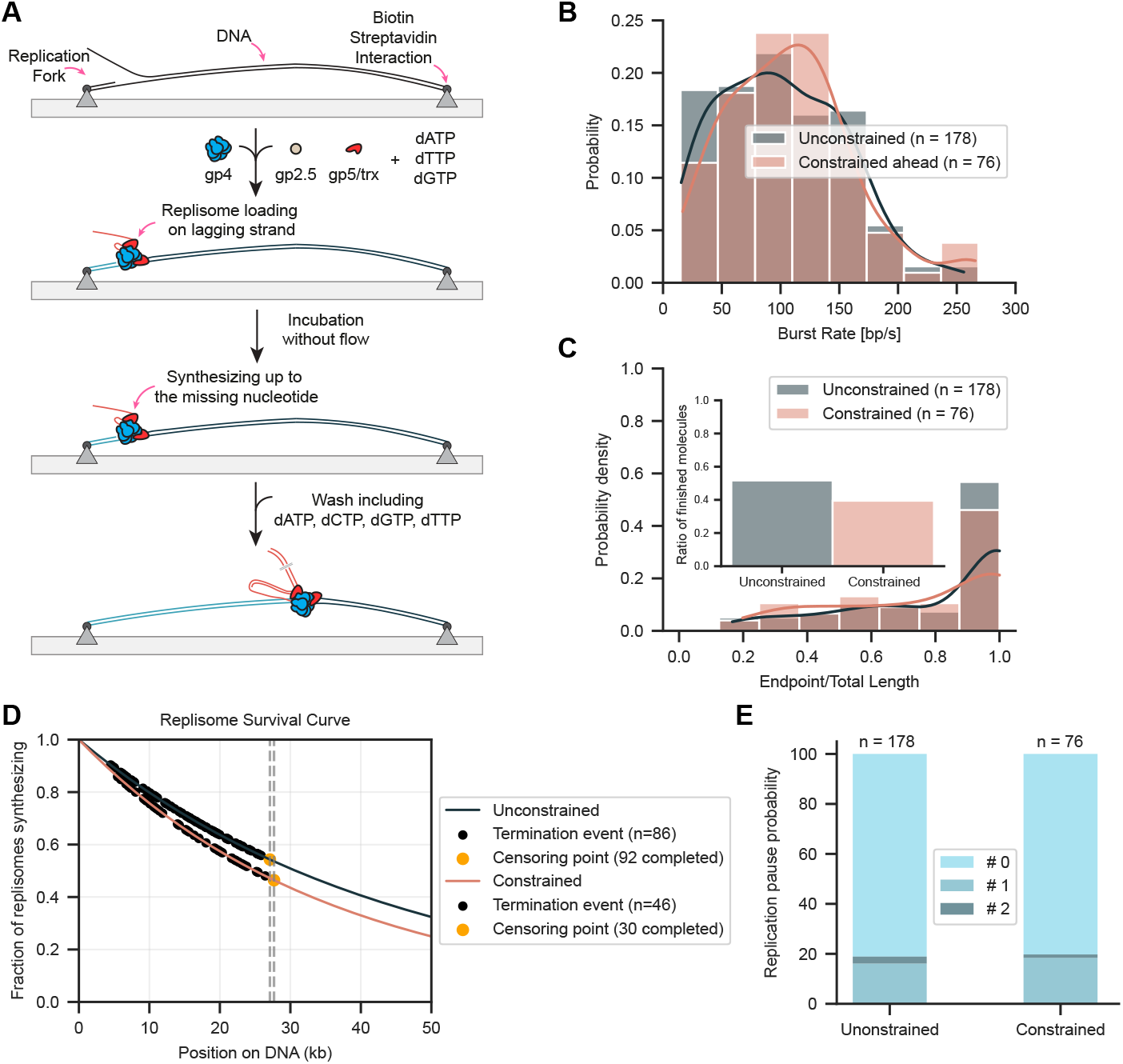
Replication using pre-assembled replisomes. (**A**) Schematic of replication assay using pre-assembly condition. (**B**) Replication burst rate distribution for unconstrained and constrained molecules. (**C**) Replication processivity distribution for unconstrained and constrained molecules. Inset displays the fraction of molecules that replicated to the end. (**D**) Replication processivity estimation using maximum likelihood estimation (**E**) Pause probabilities for unconstrained and constrained molecules.

**Figure S17.**
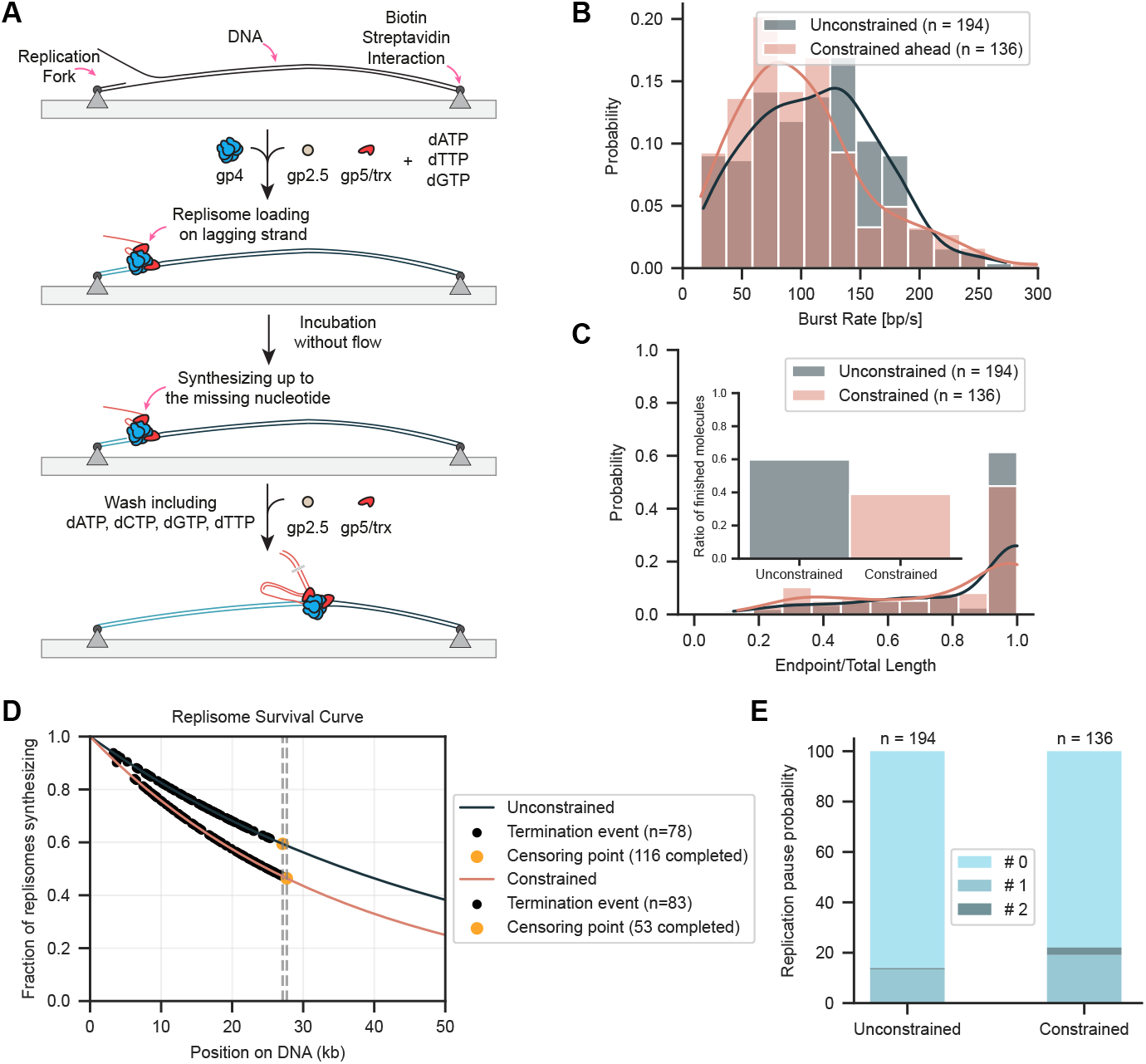
Pre-assembly condition with additional gp5 during replication. Schematic of replication assay under pre-assembly conditions with additional gp5 during replication. (**B**) Replication burst rate distribution for unconstrained and constrained molecules. (**C**) Replication processivity distribution for unconstrained and constrained molecules. Inset displays the fraction of molecules that replicated to the end. (**D**) Replication processivity estimation using maximum likelihood estimation. (**E**) Pause probabilities for unconstrained and constrained molecules.

## Supplementary Videos

**Movie S1.**
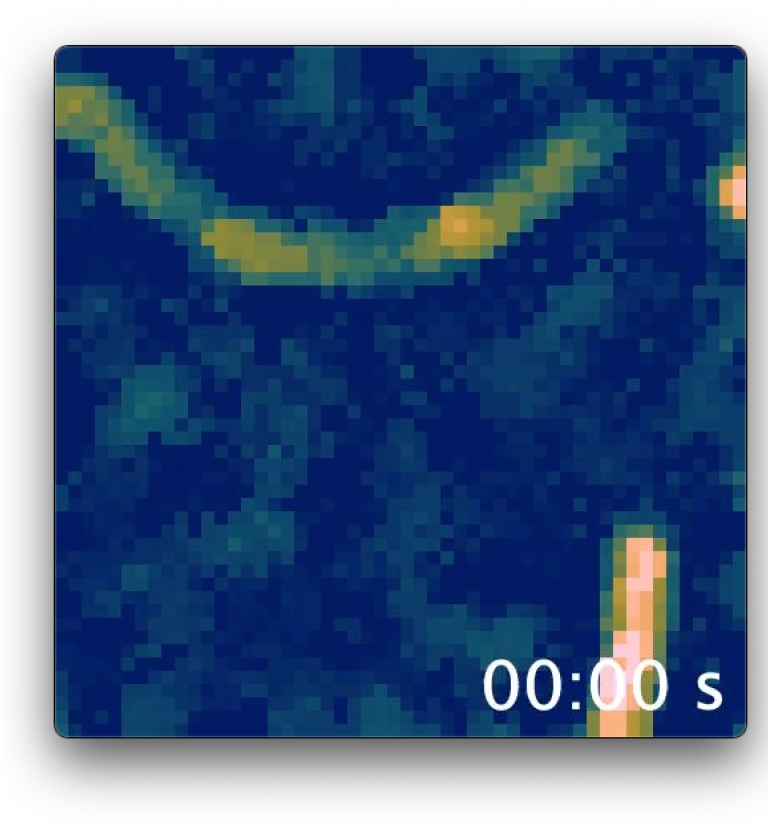
Representative unconstrained molecule during DNA replication imaged using transverse flow. The individual leading, lagging, and parental strands are all spatially resolved. The lagging-strand product is seen growing out from along the arch and extending downward due to applied flow. Intensity is color-coded using the Batlow LUT. Time is displayed as minutes and seconds in the format (mm:ss).

**Movie S2.**
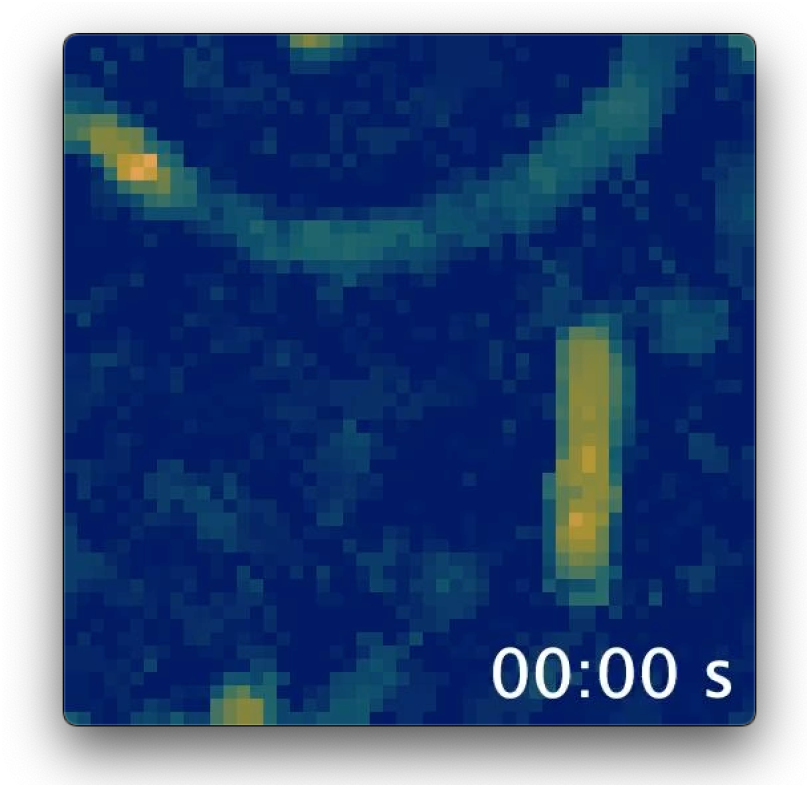
Representative constrained molecule during DNA replication imaged using transverse flow. The lagging-strand product is seen wrapping around the arch during ongoing DNA synthesis due to fork rotation. Transient downward extension events are observed in between bursts of fork rotation. Intensity is colorcoded using the Batlow LUT. Time is displayed as minutes and seconds in the format (mm:ss).

## Supplementary Tables

**Table S1.** Summary statistics from single-molecule microscopy experiments organized by experimental conditions and assay modification, reporting rates, processivities, fraction fully replicated, blob size analysis, number of polymerases and molecule numbers.

| Experimental Type | Modification | DNA Type | Mean Burst Rate [bp/s]<br>Mean±SEM | Processivity Estimation [kb]<br>Mean (95% CI) | Fraction of Fully replicated molecules [%] | Number of molecules | Figure Ref. |
| --- | --- | --- | --- | --- | --- | --- | --- |
| Linear |  | Uncon. | 99 ± 3 | 117 (102, 137) | 75 | 186 | 2 |
|  |  | Con. | 96 ± 5 | 52 (43, 67) | 57 | 79 | 2 |
|  | Pre-assembly | Uncon. | 101 ± 3 | 44 (39,52) | 52 | 178 | S16 |
|  | Pre-assembly | Con. | 108 ± 5 | 36 (29, 47) | 39 | 76 | S16 |
|  | Pre-assembly+gp5 | Uncon. | 112 ± 3 | 52 (46, 61) | 60 | 194 | S17 |
|  | Pre-assembly+gp5 | Con. | 100 ± 4 | 36 (31, 43) | 39 | 136 | S17 |
|  | Gyrase (5nM) | Con. | 103 ± 5 | 84 (70, 105) | 71 | 97 | S5 |
| Transverse |  | Unconstrained (leading) | 97 ± 7 | 84 (60, 140) | 67 | 24 | 3, 4, S7 |
|  |  | Constrained (leading) | 98 ± 9 | 29 (21, 46) | 32 | 28 | 4, S7 |
| Pausing |  |  |  |  |  |  |  |
| Experimental Type | Modification | DNA Type | 0 Pause | 1 Pause | 2 Pause | Number of molecules |  |
| Linear |  | Uncon. | 0.88 | 0.12 | 0.01 | 186 | 2 |
|  |  | Con. | 0.78 | 0.18 | 0.04 | 79 | 2 |
|  | Pre-assembly | Uncon. | 0.81 | 0.16 | 0.03 | 178 | S16 |
|  | Pre-assembly | Con. | 0.80 | 0.18 | 0.01 | 76 | S16 |
|  | Pre-assembly+gp5 | Uncon. | 0.86 | 0.13 | 0.01 | 194 | S17 |
|  | Pre-assembly+gp5 | Con. | 0.78 | 0.18 | 0.03 | 79 | S17 |
|  | Gyrase (5nM) | Uncon. | 0.9 | 0.09 | 0.01 | 97 | S5 |
| Blob size |  |  |  |  |  |  |  |
| Experimental Type |  | DNA Type | Blob size [kb] Mean±SD |  |  | Number of molecules |  |
| Linear |  | Uncon. | 1.1 ± 0.2 |  |  | 114 | 2 |
|  |  | Con. | 1.5 ± 0.4 |  |  | 52 | 2 |
| Number of Polymerases |  |  |  |  |  |  |  |
| Experimental Type | Modification | DNA Type | Mean number of polymerases Mean±SEM |  |  | Number of molecules |  |
| Linear |  | Uncon. | 3.75 ± 0.08 |  |  | 76 | 5 |
|  |  | Con | 11.40 ± 0.22 |  |  | 55 | 5 |
|  | Pre-assembly | Uncon. | 2.07 ± 0.04 |  |  | 93 | 5 |
|  | Pre-assembly | Con | 2.92 ± 0.08 |  |  | 55 | 5 |
| Linear | Pre-assembly+gp5 | Uncon. | 3.61 ± 0.05 |  |  | 165 | 5 |
|  | Pre-assembly+gp5 | Con | 4.91 ± 0.09 |  |  | 97 | 5 |

**Table S2.**
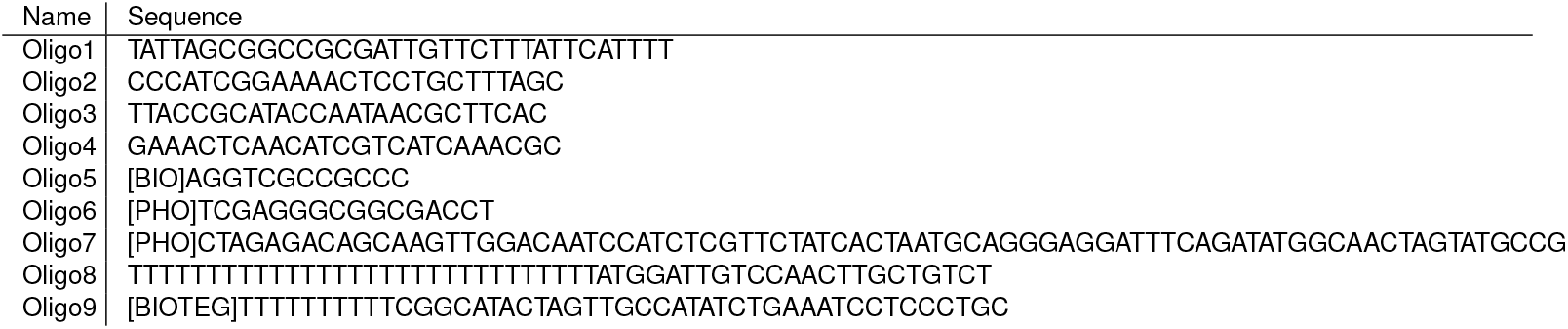
List of sequences used to create replication substrates for single-molecule experiments.

